# Post-Saccadic Disruption of Semantic Category Information in Naturalistic Scenes

**DOI:** 10.1101/2025.06.06.658316

**Authors:** Yong Min Choi, Tzu-Yao Chiu, Julie D. Golomb

**Author notes:** Corresponding author information Postal address: Moore Hall, 3 Maynard St, Hanover, NH 03755, Email address.

## Abstract

During natural vision, people make saccades to efficiently sample visual information from complex scenes. However, a substantial body of evidence has shown impaired visual information processing around the time of a saccade. It remains unclear how saccades affect the processing of high-level visual attributes – such as semantic category information–which are essential for navigating dynamic environments and supporting complex behavioral goals. Here, we investigated whether/how the processing of semantic category information in naturalistic scenes is altered immediately after a saccade. Through both human behavioral and neuroimaging studies, we compared semantic category judgments (Experiments 1A and 1B) and neural representations (Experiment 2) for scene images presented at different time points following saccadic eye movements. In the behavioral experiments, we found a robust reduction in scene categorization accuracy when the scene image was presented within 50 ms after saccade completion. In the neuroimaging experiment, we examined neural correlates of semantic category information in the visual system using fMRI multivoxel pattern analysis (MVPA). We found that scene category representations embedded in the neural activity patterns of the parahippocampal place area (PPA) were degraded for images presented with a short (0–100 ms) compared to a long post-saccadic delay (400–600 ms), despite no corresponding reduction in overall activation levels. Together, these findings reveal that post-saccadic disruption extends beyond basic visual features to high-level visual attributes of naturalistic scenes, highlighting a limitation of visual information processing in the short post-saccadic period before executing the next saccade.

## Introduction

When viewing complex visual scenes, people make saccadic eye movements - rapid, ballistic shifts of fixation to different spatial locations – to efficiently sample visual information (Najemnik & Geisler, 2005; Rayner, 2009; Renninger et al., 2007; Samonds et al., 2018; Yarbus, 1967). In many cases, saccades are functionally beneficial, projecting relevant information onto the retinal region with the highest spatial resolution, and maximizing information gain while reducing perceptual uncertainty (Renninger et al., 2007).

Although people are often unaware of any instability, numerous studies have documented various perceptual changes around the time of saccadic eye movements. During a saccade, visual input is transiently suppressed–a phenomenon known as a saccadic suppression–which supports the conscious impression of seamless stability by minimizing retinal blur (Benedetto & Morrone, 2017; Burr et al., 1994; Idrees et al., 2020; Kleiser et al., 2004). Beyond suppression effects, perception can also be distorted shortly before and after saccadic eye movements. For instance, saccades introduce perisaccadic compression, a transient distortion in spatial geometry that causes systematic mislocalization of visual objects flashed shortly before saccade onset (Ross et al., 1997; Hamker et al., 2008). Saccades also introduce perceptual instability. Because a saccade results in a displacement of retinal input, retinotopic neural representations must be remapped with each saccade (Duhamel et al., 1992; Neupane et al., 2020; Zirnsak & Moore, 2014), and this process is not perfectly efficient or instantaneous (Golomb & Mazer, 2021; Golomb & Kanwisher, 2012b). Thus, the processing of objects presented immediately after a saccade can be disrupted, including increased feature interference from objects in different locations (Golomb et al., 2014; Dowd & Golomb, 2020). If individual objects and features can be distorted immediately following a saccade, what might that mean for perception of rich, naturalistic scenes? Here we focus specifically on errors of post-saccadic perception because each saccade introduces new visual input to the retina, requiring the visual system to process novel visual information within a few hundred milliseconds before executing the next saccade.

Despite existing evidence of altered low-level perception following saccadic eye movements, it remains unclear how saccadic eye movements impact the encoding of more abstract visual information critical for everyday behavior. Visual scenes are extremely complex, in a way that multiple layers of low- and high-level visual properties are spatially organized with redundancy and regularity to form a meaningful scene (Geisler, 2008; Kersten, 1987; Malcolm et al., 2016). While some studies have used naturalistic stimuli to examine the perceptual consequences of saccades, they have largely focused on low-level properties such as local contrast (Dorr & Bex, 2013) or spatial frequency content of scene images (Kwak et al., 2024). However, successful behavior in complex visual environments requires encoding both low-level *and* high-level attributes–such as semantic category, navigability, action affordance, etc. (see Malcolm et al., 2016 for review).

How might saccadic eye movements influence the subsequent encoding of semantic category information (e.g., mountain, city, highway, etc.) from naturalistic scene images? One possibility is that the processing of semantic category information may be resilient to post-saccadic interference due to the redundant visual cues in natural scenes (Geisler, 2008; Kersten, 1987; Võ et al., 2019). Because semantic category information could be extracted from either basic-level (Castelhano & Henderson, 2008; Oliva & Schyns, 2000; Walther & Shen, 2014) or complex visual properties, such as spatial layout (Ross and Oliva, 2011) or global summary statistics (Greene & Oliva, 2009; Oliva & Torralba, 2006), previous findings on basic-level visual features may not be readily generalized to the semantic category information in naturalistic scene images. Alternatively, considering the linkage between processing of basic visual features and semantic category information (Groen et al., 2013; 2017), semantic category representations may be disrupted post-saccadically analogously to the processing of basic-level visual features.

Another intriguing alternative is that post-saccadic disruptions of semantic category representations may be more nuanced, perhaps depending on the spatial frequency conveying the scene contents. A prominent theory of rapid scene perception, the Coarse-to-Fine (CtF) model, suggests distinct roles of low and high spatial frequencies (Hegdé, 2008; Schyns & Oliva, 1994): The low spatial frequency (LSF) information conveys an abstract and coarse summary of a scene image (e.g., global layout) through the rapid magnocellular pathway, while the high spatial frequency (HSF) information carries finer details of a scene image (e.g., object details) through the relatively slower parvocellular pathway (Kauffmann et al., 2014). Given a short post-saccadic period to process the full-spectrum of spatial frequency information, the visual system may preferentially use LSF to encode scene attributes, resulting in the processing of HSF visual information being more vulnerable to post-saccadic disruption.

The current study examined whether, and how, the processing of semantic category information in naturalistic scenes is disrupted immediately after saccadic eye movements. We addressed this question using complementary behavioral and neuroimaging approaches. First, we tested behavioral performance on an explicit semantic categorization task for scene images (sampled from beach, city, forest, highway, mountain, and office categories) presented at varying time points after a saccade. Next, to assess underlying neural encoding of semantic category information in an orthogonal task, we examined the neural representation of semantic categories using functional Magnetic Resonance Imaging (fMRI) combined with multi-voxel pattern analysis (MVPA). In both cases, to examine whether the influence of post-saccadic delay is modulated by the spatial frequency conveying scene content, scene images were filtered with different spatial frequency filters to contain either low or high spatial frequency information. Previewing the findings, both behavioral and neural results indicated that the processing of semantic scene category information is transiently impaired when images are presented shortly after a saccade. These findings highlight a fundamental tradeoff in active scene perception: while saccades serve a functional benefit by projecting relevant information onto the retinal region with the highest acuity, they can also incur brief consequences for perception.

## Results

### Experiment 1

Human participants performed a gaze-contingent behavioral task in which they made a guided saccade and then reported the category of a naturalistic scene image presented after the saccade (Figure 1A). To examine how scene categorization performance varies over the post-saccadic period, we manipulated the delay between saccade offset and scene onset (Post-saccadic delay condition). In Experiment 1A, scenes appeared either 5 ms or 500 ms after the saccade. In Experiment 1B, five logarithmically spaced delays were used (5, 16, 50, 158, and 500 ms) to capture finer-grained temporal dynamics. Additionally, to test whether spatial frequency modulates the effect of post-saccadic delays on scene categorization performance, we presented scene images filtered to contain either full-spectrum (FS), high-spatial frequency (HSF), or low-spatial frequency (LSF; Figure 1B). The FS condition trials were only used to gauge overall performance and for excluding subjects.

**Figure 1.**
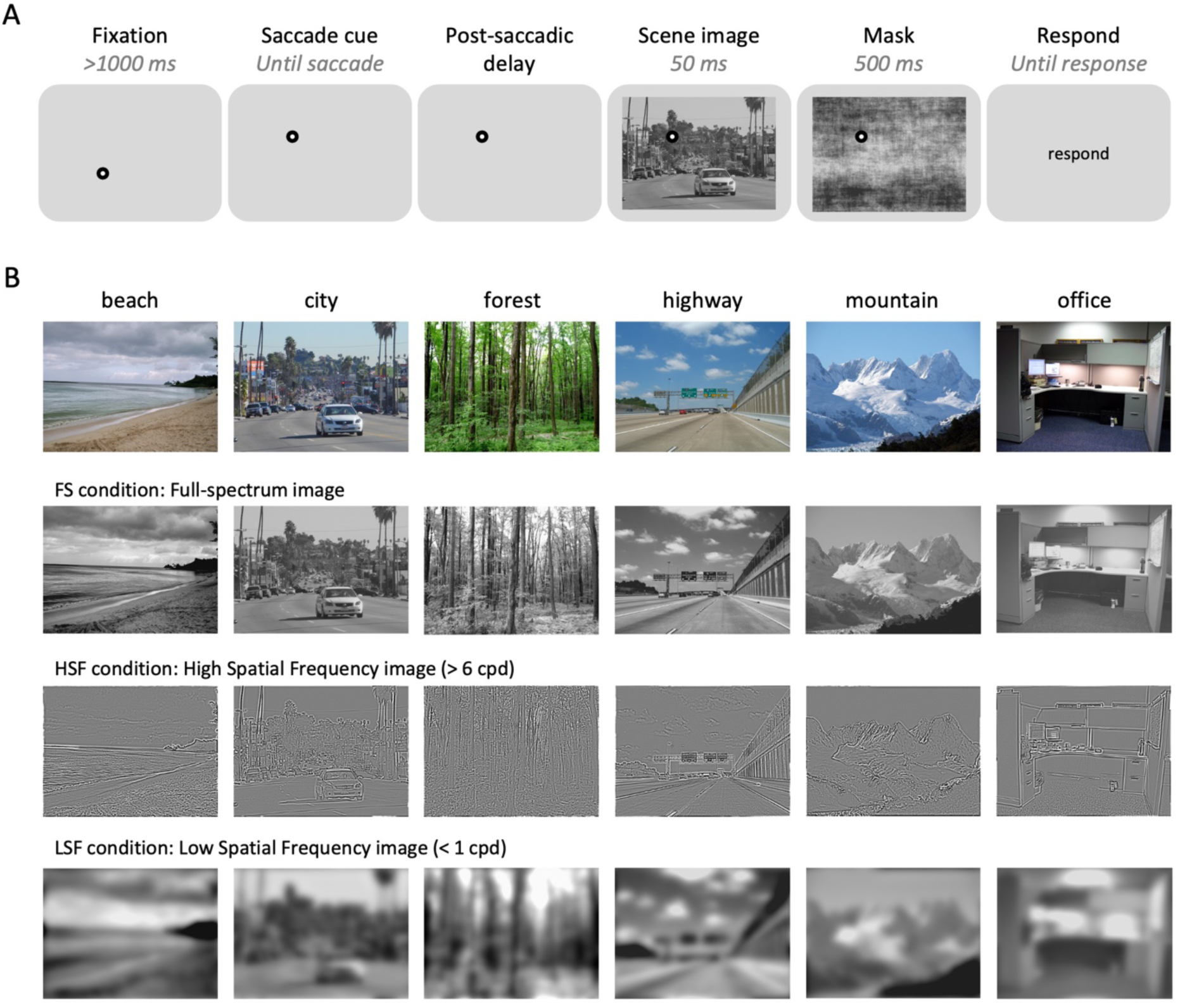
Experiment 1 design. (A) Trial sequence for behavioral experiments (Experiments 1A and 1B). (B) Example scene images of different scene categories, original images in the top row, followed by scene images filtered with different spatial frequency filters.

#### Experiment 1A

Scene categorization accuracy exceeded chance level (0.16) for both HSF (mean = 0.59, sd = 0.12) and LSF (mean = 0.62, sd = 0.12) conditions. We compared accuracies between the two post-saccadic delay conditions (5 ms vs. 500 ms) and two SF conditions (HSF vs. LSF) by performing 2 × 2 repeated-measures ANOVA (Figure 2A). We found a significant main effect of post-saccadic delay (*F*(1,20) = 17.11, *p* < .001, 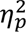 = .46, *BF*_incl_ = 189.06), with lower categorization accuracy in the 5 ms compared to the 500 ms post-saccadic delay condition. However, we found no significant main effect of the SF condition (*F*(1,20) = 2.11, *p* = .162, 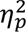 = .09, *BF*_incl_ = 0.76), nor significant interaction effect between the delay and SF condition (*F*(1,20) = 0.07, *p* = .790, 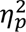 = .004, *BF*_incl_ = 0.31). These findings suggest that the processing of semantic category information is disrupted when a scene image is presented briefly following a saccadic eye movement, regardless of spatial frequency conveying the scene content.

**Figure 2.**
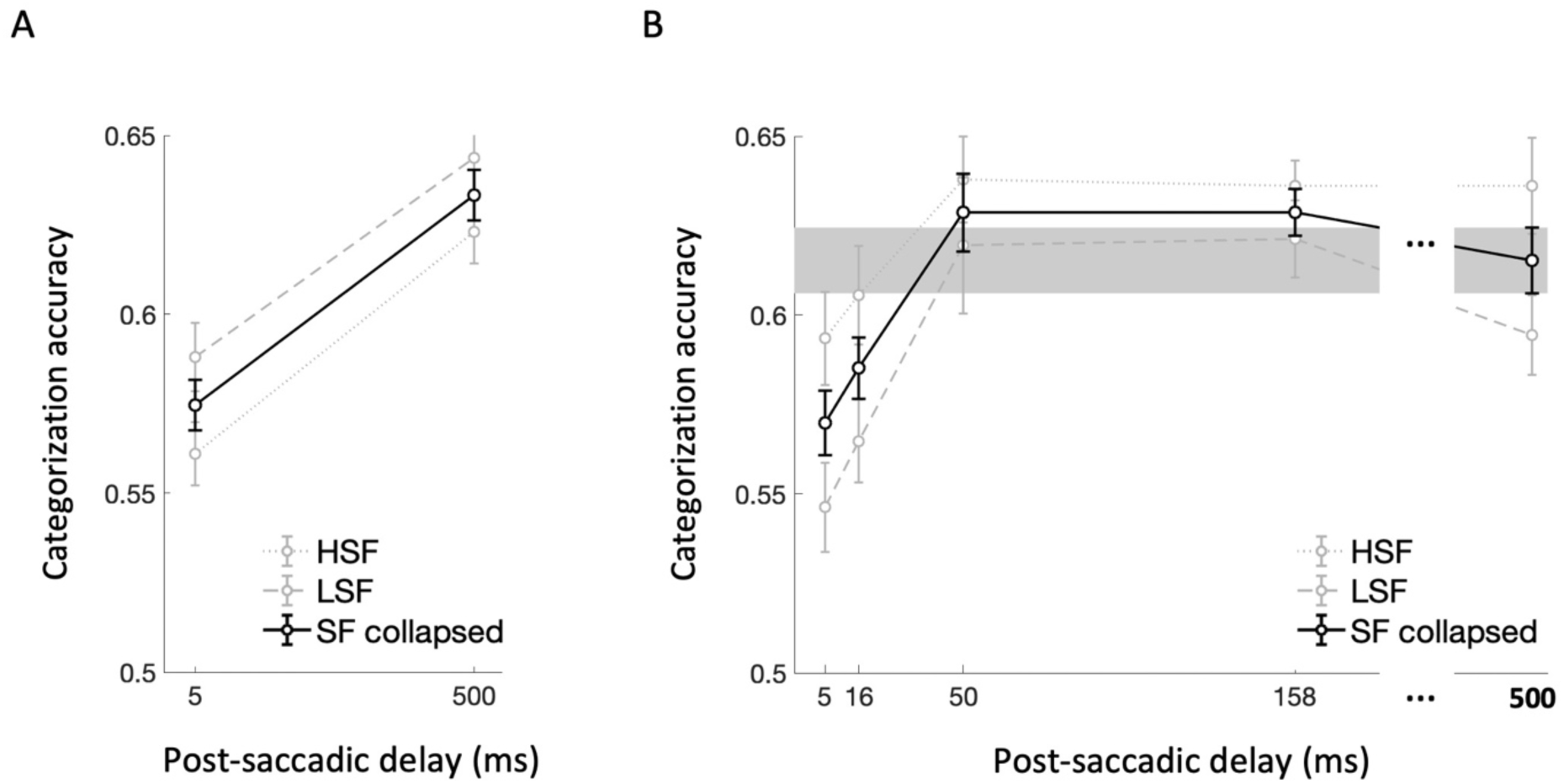
Scene categorization task results for (A) Experiment 1A and (B) Experiment 1B. Scene categorization accuracy was compared across two post-saccadic delay conditions in Experiment 1A and five post-saccadic delay conditions in Experiment 1B. In both figure, faint gray lines indicate categorization accuracy for LSF (coarse dots) and HSF (fine dots) conditions, while the solid black line indicate the scene categorization accuracy collapsed across SF conditions. For Experiment 1B, categorization accuracy in the 500 ms post-saccadic delay was used as a baseline and depicted with the gray region representing standard error. Error bars indicate within-subject standard errors. NOTE: Chance level is 0.16 for 6-AFC scene categorization task.

#### Experiment 1B

Scene categorization accuracy pooled over delay condition exceeded chance level (0.16) for both HSF (mean = 0.62, sd = 0.10) and LSF (mean = 0.59, sd = 0.12) conditions. The 5 (post-saccadic delay condition) × 2 (SF condition) repeated-measures ANOVA (Figure 2B) revealed a significant main effect of the post-saccadic delay condition (*F*(4,68) = 7.15, *p* < .001, 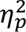 = .30, *BF*_incl_ = 75.54), without significant interaction effect with spatial frequency (*F*(4,68) = 0.53, *p* = .713, 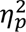 = .03, *BF*_incl_ = 0.07), consistent with Experiment 1A. While the Bayesian evidence supported a main effect of spatial frequency condition (*BF*_incl_ = 18.25), it did not reach significance with the frequentist approach (*F*(1,17) = 3.78, *p* = .069, 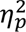 = .18).

As pre-registered, we then conducted post-hoc *t*-tests after collapsing the spatial frequency condition (Figure 2B, solid black line). Specifically, scene categorization accuracy in the 500 ms post-saccadic delay condition was considered as the baseline for recovered performance (Figure 2B, gray region) and compared with the other shorter post-saccadic delay conditions (5, 16, 50, 158 ms). We found significantly lower categorization accuracy in the 5 ms (*t*(17) = −2.87, *p* = .011, *d* = −0.68, *BF*_10_ = 5.04) compared to the 500ms baseline. Additionally, though it did not reach significance based on corrected alpha value (.0125), scene categorization accuracy was also lower in 16 ms post-saccadic delay conditions compared to the baseline (*t*(17) = −2.71, *p* = .015, *d* = −0.64, *BF*_10_ = 3.80). However, the scene categorization accuracy was not significantly different from the baseline in the 50 ms (*t*(17) = 0.73, *p* = .476, *d* = 0.17, *BF*_10_ = 0.31) and 158 ms post-saccadic delay conditions (*t*(17) = 1.29, *p* = .214, *d* = 0.30, *BF*_10_ = 0.50). Combined, these results demonstrated the time course of semantic category representation in the post-saccadic period, characterized by a significant drop in scene categorization performance shortly following the saccade offset and rapid recovery back to the baseline within 50 ms after the saccade offset.

#### Experiment 1B Exploratory analyses

For the above analyses we defined saccade offset in a real-time gaze-contingent manner, as the time when the distance between the current gaze location and the saccade target location becomes smaller than 2°. While this method is commonly used in literature, it likely underestimates saccade offset time, such that the eye may still be moving for a brief period of time after this marker. Indeed, when we performed post-hoc analyses calculating eye movement velocity at different time points relative to the scene onset time, eye movement velocity at scene onset was higher with short post-saccadic delays (Figure 3A). Thus, the decreased scene categorization accuracy in shorter post-saccadic delay trials could be attributed to the residual eye movement that can smear a visual image projected to the retina.

**Figure 3.**
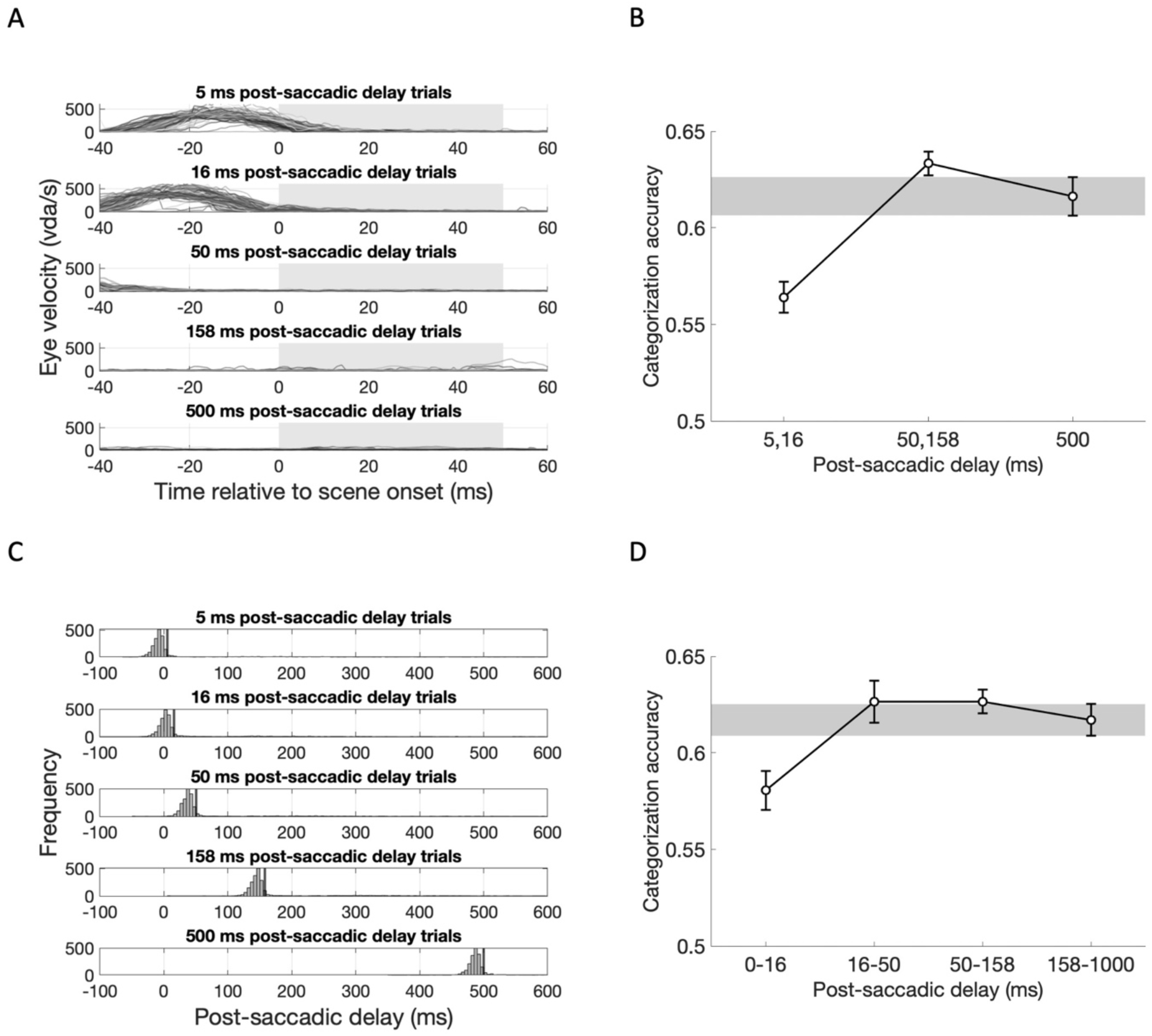
Exploratory analysis results for Experiment IB, accounting for eye movement velocity and the saccade detection algorithm. (A) Eye movement velocity (dva/sec; y-axis) as a function of time relative to scene onset (x-axis) is plotted for each post-saccadic delay condition (rows) from a single exemplar subject. Gray boxes indicate the duration of scene presentation. (B) Group-level scene categorization accuracy for short (5, 16 ms), intermediate (50, 158 ms), and long (500 ms) post saccadic delay trials, after excluding trials in which eye movement velocity exceeded 25 dva/s at scene onset. (C) Post-saccadic delay at scene onset for each condition, calculated using saccade onset times from the EyelinklOOO’s online parsing system for each subject. Black vertical lines indicate the intended post-saccadic delay. (D) Group-level scene categorization accuracy compared across post saccadic delay conditions based on re-calculated post-saccadic delays, grouped into four post-saccadic delays. Error bars indicate within-subject standard errors.

To investigate whether retinal shifts of visual input are responsible for reduced scene categorization performance, we excluded trials on which the eyes were still moving at scene onset (>25 °/sec), and compared categorization accuracy for short (5, 16 ms), intermediate (50, 158 ms) and long (500 ms) post-saccadic delay trials (Figure 3B). One-way repeated-measures ANOVA revealed a significant main effect of post-saccadic delay (*F*(2,34) = 12.99, *p* < .001, 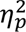 = .43, *BF*_incl_ = 305.89), characterized by significantly lower categorization accuracy for short (5, 16 ms) post-saccadic delay trials compared to intermediate (*t*(17) = −4.89, *p_bonf_* < .001, *d* = −1.15, *BF*_10_ = 6060.12) and long post-saccadic delay trials (*t*(17) = −3.69, *p_bonf_* = .002, *d* = −0.87, *BF*_10_ = 7.09), without significant difference between intermediate and long post-saccadic delay trials (*t*(17) = 0.28, *p_bonf_* = .721, *d* = 0.28, *BF*_10_ = 0.44). These results indicate that the observed post-saccadic drop in scene categorization accuracy was not due to the confound of residual eye movement.

In addition, we also employed an alternative algorithm to detect saccade onset and offset. Using the online parsing system built-in Eyelink 1000, we re-calculated trial-wise post-saccadic delay (Figure 3C). The majority of re-calculated post-saccadic delays (histograms) were shorter than the intended post-saccadic delays (black vertical lines), suggesting that this method is more strict way of defining post-saccadic delay for each stimuli onset. Then, we labeled each trial based on calculated post-saccadic delay into four post-saccadic delay groups (0-16 ms, 16-50 ms, 50-158 ms, 250-1000 ms; Figure 3D). One-way repeated-measures ANOVA again revealed a significant main effect of post-saccadic delay (*F*(3, 51) = 4.38, *p* = .008, 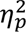 = .205, *BF*_incl_ = 5.42). Post-hoc analysis found lower scene categorization accuracy in 0-16 ms post-saccadic delay trials compared to the 16-50 ms (*t*(17) = −3.12, *p*_bonf_ = .019, *d* = −0.73, *BF*_10,U_ = 3.34) and 50-158 ms (*t*(17) = −3.12, *p*_bonf_ = .018, *d* = −0.74, *BF*_10,U_ =8.64), with marginal difference compared to the 158-1000 ms post-saccadic delay trials (*t*(17) = −2.47, *p*_bonf_ = .10, *d* = −0.58, *BF*_10,U_ = 2.42). The exploratory analyses revealed impaired semantic category information for scene images presented immediately after saccadic eye movement, which is not attributed to smeared retinal image nor limited to the saccade detection methods used in the main analysis.

### Experiment 2

In Experiment 2, we adopted a neuroimaging approach, using functional Magnetic Resonance Imaging (fMRI) and multi-voxel pattern analysis (MVPA; Haxby et al, 2001), to assess whether and how neural representations of semantic scene category information are altered following saccades. Specifically, if scene content processing is disrupted post-saccadically, this should be reflected in degraded decoding of scene category information within scene-selective brain regions such as the parahippocampal place area (PPA; Epstein & Kanwisher, 1998).

Comparing neural indicators of semantic scene category information in the absence of an explicit categorization task is particularly useful to rule out alternative explanations for the reduced behavioral performance evaluated in Experiment 1. For example, non-perceptual factors such as interference with decision-making (Matsumiya & Furukawa, 2023), motor planning or execution (Pashler et al., 1993; Richardson et al., 2013) may be responsible for the reduced performance in short delay conditions and/or the absence of interaction with spatial frequency. By examining neural evidence of scene category providing direct evidence for the perceptual disruption of scene content during the post-saccadic period.

In the fMRI scanner, subjects followed a fixation dot and performed a 1-back task on sequentially presented scene images, pressing a button only when the current image was identical to the one shown on the previous trial (Figure 4A). Similar to the behavioral experiments, we aimed to compare scene images presented with either short or long delays after the saccade cue onset (Post-saccadic delay condition). To manipulate post-saccadic delay, we integrated high temporal resolution eye tracking with the fMRI system. However, unlike the fully gaze-contingent behavioral experiments, where stimulus presentation on each trial was contingent on online eye-tracking data, the fMRI trial sequence had to be pre-scheduled and time-locked to the scanner’s repetition time. Therefore, we recorded high resolution eye tracking data during each fMRI trial and used it to subsequently select trials for inclusion in which the scene onset time fell within the designated short (0-100 milliseconds) or long (400-600 milliseconds) post-saccadic delay windows. To maximize trial inclusion, we first measured each subject’s average saccade reaction time during a pre-scan session (Figure 4B), and adjusted the saccade cue onset timing for that subject in their fMRI session to maximize the likelihood that scenes would appear within these windows (see *Materials and Methods* for details). The scene image stimuli were drawn from two nature scene categories (beach and mountain) and two urban scene categories (city and highway), and were filtered with two low spatial frequency (LSF1 and LSF2) and two high spatial frequency bands (HSF1 and HSF2), matching the hierarchical structure of the scene category manipulation.

**Figure 4.**
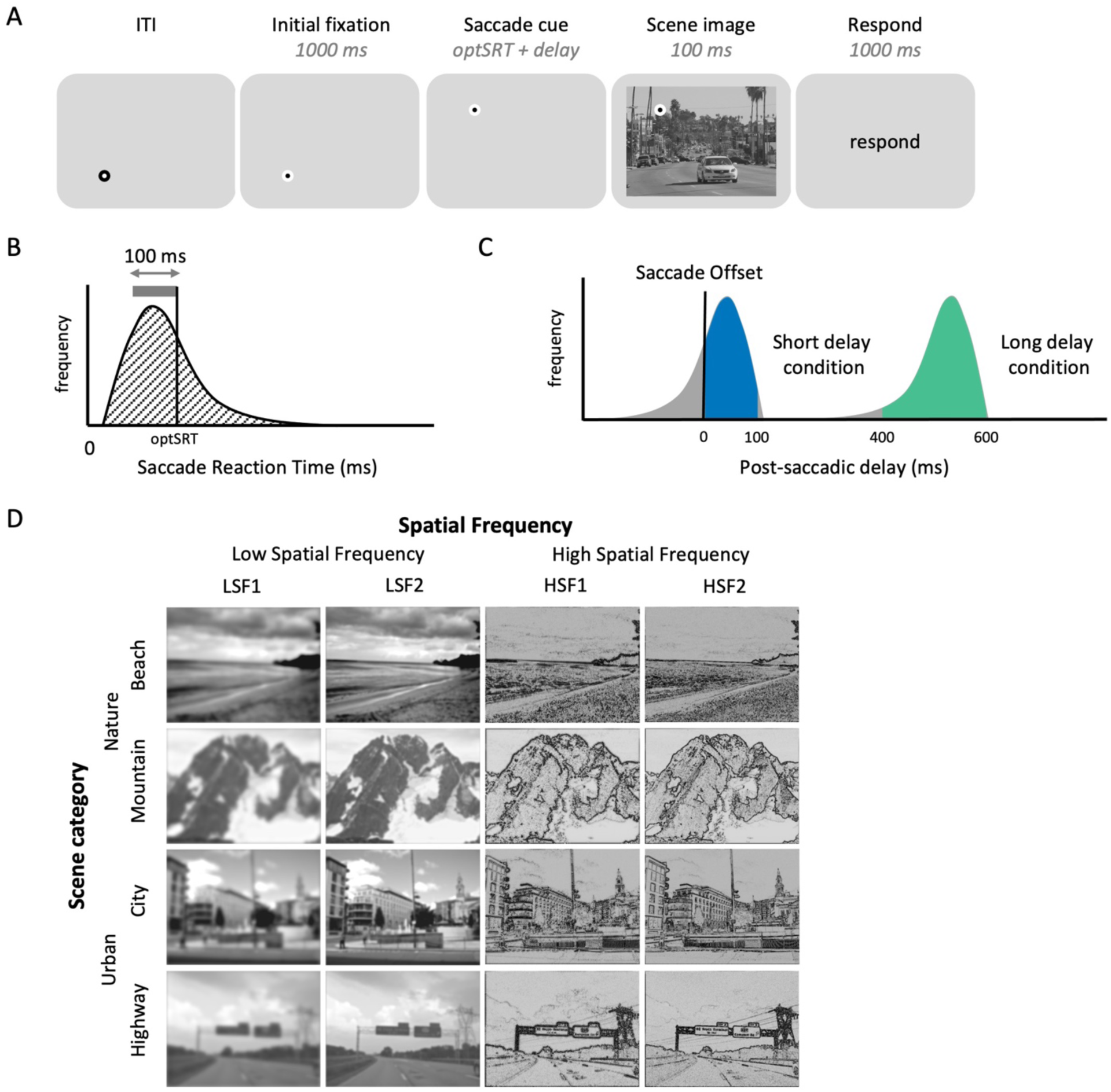
Experiment 2 design. (A) Trial sequence. (B) illustrative distribution of saccade reaction time (ms) - delay between saccade cue onset to saccade offset - measured during the pre-scan session. We identified a 100 ms range containing the highest concentration of saccade reaction times (gray horizontal bar). The upper bound of this range (black vertical bar) was defined as the optimal saccade reaction times *(optSRT),* and used to set the delay between the saccade cue and scene onset in the scan session. (C) Illustrative distribution of post-saccadic delays - delay between saccade offset and scene onset - measured during the scan session. By presenting scene image either *optSRT* or *optSRT* + *500 ms* after the saccade cue onset, trial-wise post-saccadic delays followed a bimodal distribution, maximizing the number of short (0-100 ms; blue) or long (400-600 ms; green) post-saccadic delay trials, selected through post-hoc analyses of eye-tracking data and included in the main analysis. (D) Each scene image was labeled as one of four scene category conditions and four spatial frequency conditions.

#### Post-saccadic disruption of scene category information in PPA

To quantify the amount of scene category information (natural vs. urban) represented in PPA, we performed a multi-voxel pattern analysis (MVPA; Haxby et al., 2001; Golomb & Kanwisher, 2012a). Specifically, we constructed representational similarity matrices (RSMs) and tested whether voxel-wise activation patterns in PPA were more similar between trials featuring the same scene category than between trials with different categories (Figure 5; see *MVPA analysis* section for details).

We found that activity patterns were more similar for images of the same scene category than for different categories, across all spatial frequency and post-saccadic delay conditions (*p*s<.021, *BF*_10_s >2.97; Figure 6A), indicating significant semantic scene category representation encoded in PPA. Next, to assess how scene category information (indexed by the same-minus-different category difference scores) was influenced by post-saccadic delay and its interaction with spatial frequency, we calculated scene category representation on 8 × 8 RSMs (correlations within each delay x SF condition) and conducted a 2 (Post-saccadic delay condition) × 2 (Spatial frequency condition) repeated measures ANOVA (Figure 6B). The ANOVA showed no significant interaction (*F*(1,16) = 1.622, *p* = .221, 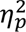 = .09, *BF*_incl_ = 0.61) or main effect of spatial frequency (*F*(1,16) = 3.38, *p* = .085, 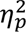 = .17, *BF*_incl_ = 2.11). Nevertheless, there was significant main effect of post-saccadic delay condition (*F*(1,16) = 9.99, *p* = .006, 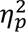 = .38, *BF*_incl_ = 1.07), indicated by reduced scene category information in the short compared to the long post-saccadic delay trials. A post-hoc analysis of simple main effects revealed reduced scene category information in short compared to long post-saccadic delay trials in the HSF condition (*F*(1) = 6.94, *p* = .018, *d* = −0.64, *BF*_10_ = 3.31), but not in the LSF condition (*F*(1) = 0.26, *p* = .615, *d* = −0.12, *BF*_10_ = 0.28).

**Figure 6.**
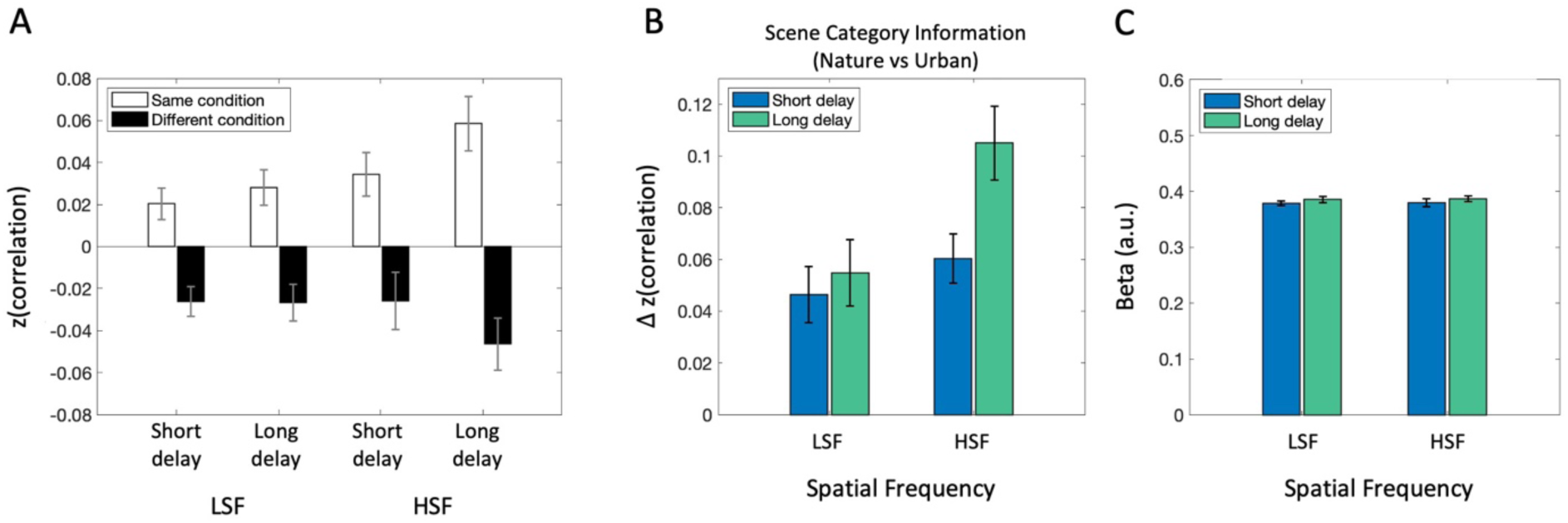
fMRI analysis results in PPA. (A) Average correlation between the same scene category pairs (white bars) and different category pairs (black bars) shows more similar neural activity patterns between same scene category trials than different scene category trials. (B) Scene category representation in PPA, obtained by subtracting average correlation between different scene category pairs from same category pairs, separately for spatial frequency conditions (LSF vs. HSF) and post-saccadic delay conditions (Short vs. Long). (C) Univariate analysis result showing overall activation level between each condition. Error bars indicate within-subject standard error.

Next, to better capture the broader effect of post-saccadic delay, we performed another MVPA analysis separating the RSM only by delay (16 × 16 cell RSMs) to calculate scene category information at each delay regardless of spatial frequency. A paired-samples *t*-test confirmed significantly lower scene category information in the short post-saccadic delay trials (*M* = 0.053) compared to that of long post-saccadic delay trials (*M* = 0.078; *t*(16) = −3.84, *p* = .001, *d* = −0.93, *BF*_10_ = 27.64).

#### Disrupted neural activity pattern without reduced activation

Is the reduction in semantic category representation in neural activity pattern driven by an overall reduction in PPA activation to scene images? We conducted a standard univariate analysis averaging beta estimates in PPA (Figure 6C). A 2 (Post-saccadic delay condition) × 2 (Spatial frequency condition) repeated measures ANOVA revealed no significant main effect of post-saccadic delay (*F*(1,16) = 0.86, *p* = .366, 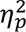 = .051, *BF*_incl_ = 0.41), nor a significant interaction (*F*(1,16) = .001, *p* = .974, 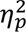 = .00, *BF*_incl_ = 0.31) or main effect of spatial frequency (*F*(1,16) = 0.023, *p* = .881, 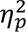 = .001, *BF*_incl_ = 0.25). The absence of post-saccadic delay effect on univariate activation suggests that overall activation to visual scene stimuli remains intact after a saccade, even when the neural activation pattern encoding the semantic scene content was disrupted.

#### Consistent patterns in PPA subregions along anterior-posterior axis

Motivated by the functional distinction of PPA along the anterior-posterior axis (Baldassano et al., 2016; Berman et al., 2017), we further investigated if sub-regions of PPA along the anterior-posterior axis are differently influenced post-saccade. We performed a 2 (Post-saccadic delay condition) × 2 (Spatial frequency condition) × 2 (PPA subregions) repeated measures ANOVA (Figure 7A). Consistent with post-saccadic disruption of scene category information in PPA as a whole, we found a main effect of post-saccadic delay on scene category information (*F*(1,16) = 7.17, *p* = .016, 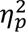 = .31, *BF*_incl_ = 1.37), without significant interaction between post-saccadic delay and spatial frequency (*F*(1,16) = 1.50, *p* = .238, 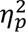 = .09, *BF*_incl_ = 1.35). Moreover, there was a significant higher scene category information in HSF, compared to the LSF condition (*F*(1,16) = 4.772, *p* = .044, 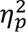 = .23, *BF*_incl_ = 32.05). Importantly, we did not find any 2-way nor 3-way interaction effects involving PPA subregions (*p*s>0.67, *BF*_10_s < .24), suggesting no functional distinction between anterior and posterior PPA concerning the post-saccadic processing of semantic category information. Consistent with the univariate results for the PPA overall, a 2 × 2 × 2 repeated measures ANOVA on univariate activation (Figure 7B) did not find any significant main effects nor interactions with subregion (*p*s > .40, *BF*_incl_s < 0.46).

**Figure 7.**
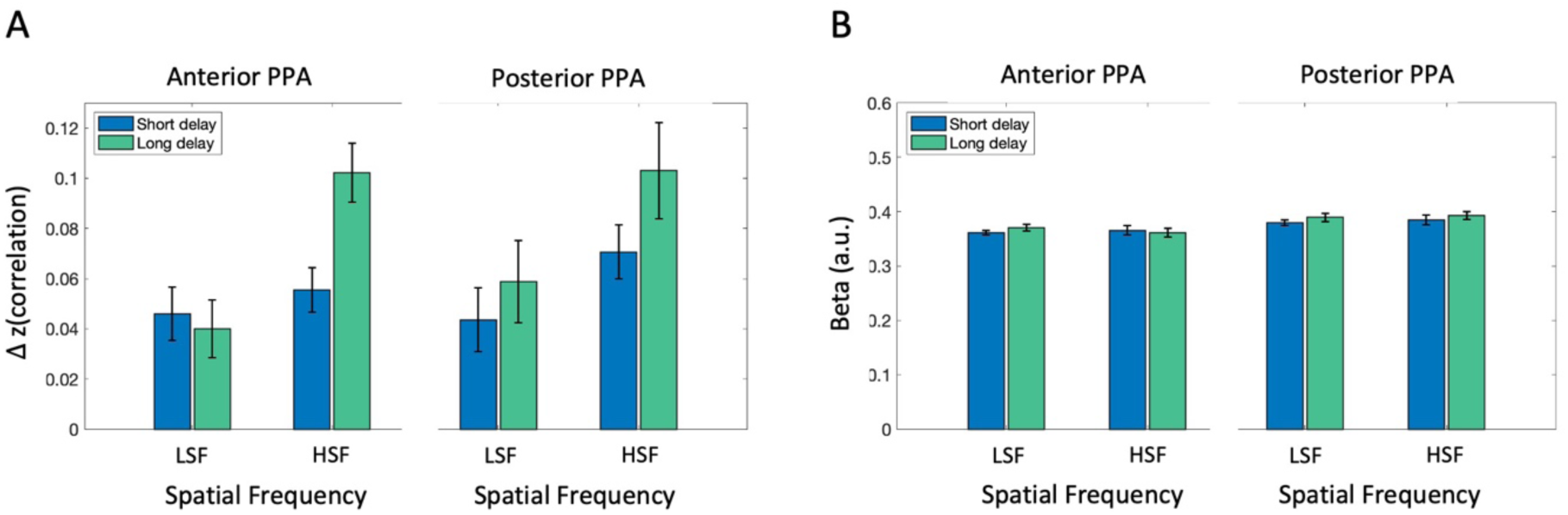
fMRI analysis results in anterior and posterior PPA (A) Scene category representation in anterior (left) and posterior PPA (posterior), obtained by subtracting average correlation between different scene category pairs from same category pairs, separately for spatial frequency conditions (LSF vs. HSF) and post-saccadic delay conditions (Short vs. Long). (C) Univariate analysis result showing overall activation level in each condition, separately for anterior (left) and posterior PPA (right). Error bars indicate within-subject standard error.

### Spatial frequency information in early visual cortex

Finally, while our primary focus is on post-saccadic representations of semantic scene content (scene category information), the spatial frequency manipulation also allowed us to examine post-saccadic processing of basic-level visual features in complex scene images (i.e., spatial frequency information). Similar to above, we conducted both MVPA and univariate analyses, now examining the amount of spatial frequency information (LSF vs. HSF) in the early visual cortex (EVC). As shown in Figure 8A, the MVPA analysis tested whether there was significant information in the pattern of EVC response to differentiate whether a scene contained high vs low spatial frequency content. EVC exhibited significant scene frequency information in both short (*t*(16) = 3.69, *p* = .002, *d* = 0.90, *BF*_10_ = 21.126) and long post-saccadic delay trials (*t*(16) = 2.58, *p* = .02, *d* = 0.63, *BF*_10_ = 3.04). Although the magnitude was numerically higher in the long delay, a paired-samples *t*-test revealed no significant effect of post-saccadic delay on spatial frequency representation (*t*(16) = −0.98, *p* = .34, *d* = −0.24, *BF*_10_ = 0.38). The univariate analysis also found no significant difference between the short and long post-saccadic delay condition in EVC (Figure 8B; *t*(16) = −0.92, *p* = .370, *d* = −0.224, *BF*_10_ = 0.361). Additional analyses calculating spatial frequency information within low (LSF1 vs. LSF2) or high (HSF1 vs. HSF2) spatial frequency bands were not significant in EVC (Supplementary Figure 6).

**Figure 8.**
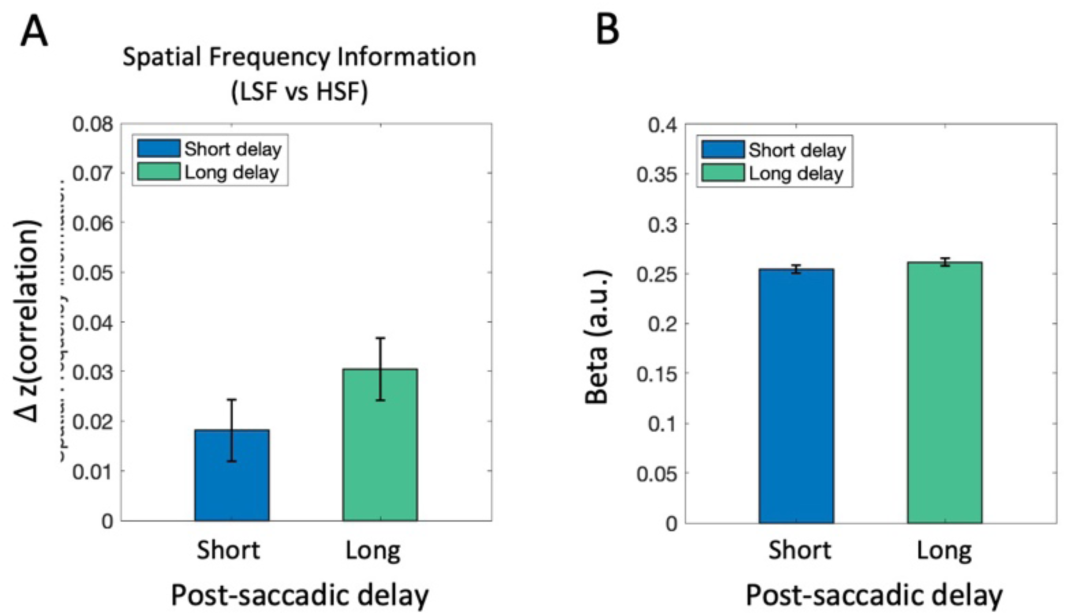
fMRI analysis results in EVC. (A) Neural representation of spatial frequency information was compared between the short and long post-saccadic delay conditions. (B) Univariate activation was compared between the short and long post-saccadic delay conditions. Error bars indicate within-subject standard error.

## General Discussion

The current study used a combination of behavioral and neuroimaging approaches to investigate an understudied aspect of naturalistic visual scene perception: whether representations of semantic scene category information are briefly altered in the time period immediately following a saccadic eye movement. Our behavioral experiments revealed significantly diminished scene categorization accuracy when the scene image was presented following the shortest post-saccadic delays (<50 ms), compared to after longer delays. Moreover, in the fMRI experiment, we assessed neural representations of semantic category in scene-selective brain region PPA using MVPA, and found analogously disrupted semantic category representations for scene images presented with short (0-100 ms) compared to longer post-saccadic delays (400-600 ms). The degraded neural representation even in absence of an explicit semantic task rules out non-perceptual explanation such as decision-making interference (Matsumiya & Furukawa, 2023) or motor planning (Pashler et al., 1993; Richardson et al., 2013), underscoring genuine disruption of semantic category representation in the post-saccadic period.

Furthermore, fMRI data revealed no effect of post-saccadic delay on univariate activation in PPA, suggesting that saccades interfere with representations of scene content (neural pattern encoding), rather than reducing overall activity. The lack of an activation difference argues against the possibility that residual eye movements restrict the amount of visual information reaching the system at early processing stages. Moreover, it may indicate that PPA still recognized the visual input as a ‘scene’, while the detailed semantic content is not fully processed post-saccadically. This proposes interesting correspondence with prior findings, where people are often surprisingly insensitive to trans-saccadic changes in scene details (Choi et al., 2025; Henderson & Hollingworth, 2003; Kwak et al., 2024), whilst maintaining a coherent conscious percept of the visual scene.

Together, these findings demonstrate that high-level visual attributes of naturalistic scene are vulnerable to disruption following saccades, despite the redundancy and regularity of naturalistic scene images (Geisler, 2008; Kersten, 1987; Malcolm et al., 2016; Võ et al., 2019). Adding to the known functional benefits of saccadic eye movements during active exploration of visual scenes, these results suggest that saccades may also carry brief costs for subsequent visual information processing.

### The effect of spatial frequency conveying semantic category information

In all experiments, we manipulated the spatial frequency content of scene stimuli to examine its influence on semantic category processing in the post-saccadic period. Particularly, inspired by the Coarse-to-Fine (CtF) model (Hegdé, 2008; Schyns & Oliva, 1994), we hypothesized that the rapid processing of high-level scene attributes may rely more on the LSF information, making HSF images more susceptible to post-saccadic disruption. On the other hand, some studies of saccadic suppression have found stronger suppression (i.e., reduced sensitivity) for LSF compared to HSF stimuli (Burr et al., 1994; Idrees et al., 2020; Kleiser et al., 2004), which would predict the opposite pattern in our study.

Our fMRI study revealed interesting effects of spatial frequency. First, we found overall stronger semantic category representations for HSF compared to LSF scene images, especially in long-post-saccadic delay trials, consistent with prior work suggesting that scene content may be predominantly conveyed by HSF information (Berman et al., 2017; Kauffmann et al., 2015; Rajimehr et al., 2011).

Interestingly, it was HSF scene images - not LSF scene images - that exhibited a significant reduction in semantic category representation in short post-saccadic delay trials, although the interaction effect was not significant. While the relative preservation of semantic information for LSF scenes under short delays is consistent with the prediction grounded on CtF model, the current findings alone are insufficient to conclude if the visual system preferentially relies on LSF information in post-saccadic scene perception. Future research could clarify how the visual system differentially processes spatial frequency information during the immediate post-saccadic period.

The greater post-saccadic impairment for HSF scene images does seem inconsistent with a stronger saccadic suppression observed with LSF compared to HSF stimuli (Burr et al., 1994; Idrees et al., 2020; Kleiser et al., 2004). This discrepancy may reflect distinctive processing of localized objects versus naturalistic scenes (Boucart et al., 2013; Hasson et al., 2002; Levy et al., 2001; Malach et al., 2002). While object recognition relies on central vision with high spatial resolution, scene processing remains robust in peripheral vision (Boucart et al., 2013) and even with low-pass filtered images (Nuthmann, 2013, 2014). Indeed, scene-selective voxels are clustered medially in the ventral temporal cortex and exhibit a preference for peripheral visual input (Grill-Spector & Weiner, 2014; Hasson et al., 2002; Levy et al., 2001; Malach et al., 2002). Taken together, the distinct patterns of post-saccadic visual perception, modulated by spatial frequency, may reflect an optimized use of different spatial frequencies around the time of saccadic eye movements for more efficient scene processing.

Unlike the fMRI experiment, the behavioral experiments did not find a corresponding effect of spatial frequency information, possibly due to insufficient sensitivity of the categorization task to capture subtle effects of low-level image statistics. The visual environment is highly complex and redundant (Geisler, 2008; Kersten, 1987; Võ et al., 2019). When explicitly categorizing scenes, observers may rely on a variety of cues - including basic features (Castelhano & Henderson, 2008; Oliva & Schyns, 2000; Walther & Shen, 2014), spatial layout (Ross & Oliva, 2011), or global summary statistics (Greene & Oliva, 2009; Oliva & Torralba, 2006) – potentially obscuring subtle effects of spatial frequency.

### The absence of disrupted spatial frequency information in EVC

Interestingly, in contrast to prior behavioral findings showing impaired sensitivity to basic-level visual features like contrast (Dorr & Bex, 2013) or spatial frequency (Kwak et al., 2024) in naturalistic scenes, our fMRI results revealed no significant effect of post-saccadic delay on the neural representation of spatial frequency information in early visual cortex. One possible explanation is that the duration of the scene image in our fMRI study (100 ms) was sufficiently long to allow adequate processing of basic-level visual information even when accounting for post-saccadic disruption. Using neuroimaging techniques with superior temporal resolution (e.g., EEG, MEG), previous studies have examined the time course of naturalistic visual stimuli processing for different attributes (Dima et al., 2018; Fakche et al., 2024).

Specifically, a recent MEG experiment showed the neural representation of object color emerging around 100 ms after saccade offset, followed by category-level information around 145 ms (Fakche et al., 2024). The more rapid processing of basic-level visual features may mean that it could have escaped from post-saccadic interference in our experiment design, particularly on trials where the scene image was presented at the later end of the short-delay window.

### Other high-level visual attributes during naturalistic scene processing

While we focused on semantic category information in naturalistic scenes, it does not capture the full range of high-level visual attributes necessary for interacting with the environment, such as action affordance and navigability (Epstein & Baker, 2019; Malcolm et al., 2016). For example, the stronger degradation of semantic category information when viewing HSF scene images may not generalize to other attributes (e.g., action affordance, navigability), considering literature suggesting distinct, flexible usage of spatial frequency information depending on task demands (Wiesman et al., 2021). Moreover, compared to some of these other attributes, semantic category is a more stable attribute over time. While the semantic category of the current visual scene generally does not change across eye movements, navigable paths—defined in egocentric coordinates—change with each fixation and must be continuously updated across saccades (Wang & Spelke, 2000; Bonner & Epstein, 2017), as do the action affordances of objects (Medendorp et al., 2008; Henrique et al., 1998; Batista et al., 1999). Future research could explore how saccades affect these more dynamic scene attributes and how the visual system interacts with motor networks to enable seamless perception and action in naturalistic environments (Goodale, 2011; Tagliabue & McIntyre, 2012).

Lastly, our findings raise a fundamental question: how do individuals navigate complex visual environments effortlessly despite disruptions in high-level visual processing after saccades? Decades of research have identified multiple mechanisms supporting trans-saccadic perceptual stability, spanning neural (Duhamel et al., 1992; Wurtz, 2008), cognitive (MacKay, 1973), and visual (Binda & Morrone, 2018) levels. While majority of these theories were built upon the stability of basic visual properties, such as spatial displacement (Deubel et al., 1996) or changes in surface features of isolated objects (Weiß et al., 2015), there is increasing recognition on testing stability mechanisms in a more ecologically valid context (Choi et al., 2025). Here, we leveraged complementary use of behavioral and neural evidence and demonstrated disrupted processing of a high-level visual attribute – semantic category information – when viewing naturalistic scene images. Our results further underscore the need for future research to explore trans-saccadic perception in naturalistic settings with dynamic task demands to fully understand how the brain achieves coherent visual experience in real-world contexts.

## Materials and Methods

### Experiment 1

#### Pre-registration Statement

Experiment 1A was not explicitly pre-registered; however, it was a modification of a similar experiment we had pre-registered (https://osf.io/az9c7), retaining the core motivation, sample size, and design. Experiment 1B was pre-registered, including its rationale, design, and analysis plan (https://osf.io/h8dmu). Any additional analyses beyond the pre-registration are reported as exploratory.

#### Participants

As pre-registered in our preliminary experiment, we set 18 subjects as a minimum sample size for Experiment 1A. This was based on a previous study (Perfetto et al., 2020) testing scene categorization performance between LSF and HSF images. We performed a Bayesian analysis on their data (Experiment 2, which reported no significant difference between conditions, *t*(17) = 0.034; *p* = 0.97) and found moderate support for the null hypothesis (*BF*_10_<.228). Thus, we planned to collect at least 18 participants and apply the Bayesian optional stopping rule (Rouder, 2014), continuing data collection by sets of three subjects to counterbalance spatial frequency condition order (see *Stimuli* section) until the Bayes factor indicated sufficient evidence either for (BF₁₀ > 3) or against (*BF*_10_<0.333) our key effect of interest: the interaction between spatial frequency (HSF vs. LSF) and post-saccadic delay (Short vs. Long). The maximum sample size was set at 36.

Ultimately, data from 21 participants (12 women, 9 men; *M* = 20.86 years, *SD* = 4.89) were included in the final analysis for Experiment 1A. Two additional participants completed the experiment but were excluded for different reasons: one due to categorization accuracy for FS images lower than preregistered exclusion criterion of 50% (49.2%), and the other due to system error that caused a longer post-saccadic delay than intended. Experiment 1B followed the same sample size plan, and data from 18 participants (13 women, 5 men; *M* = 19.00 years, *SD* = 2.37) were collected, with no additional data collection given sufficient Bayesian evidence. All participants had normal or corrected-to-normal vision and received either course credit or monetary compensation for participation ($15/hour). Experiments 1A and 1B were approved by the Ohio State University Behavioral and Social Sciences Institutional Review Board and written informed consent was obtained from all participants.

### Experiment design

#### Experiment 1A

Subjects participated in a gaze-contingent behavioral experiment, where they were instructed to follow a fixation dot with their eyes and perform a 6-AFC (i.e., beach, city, forest, highway, mountain, and office) scene categorization task on a briefly presented scene image. Each trial started with an initial fixation dot located at one corner of an imaginary 10° ×10° right square centered on the screen (Figure 1A). Once participants successfully fixated on the initial fixation for more than 1000 ms, the fixation dot disappeared and immediately reappeared at a different corner of the imaginary square (saccade cue). Subjects were instructed to make an eye movement toward the saccade cue as fast and accurately as possible. Eye position was monitored in real-time, and saccade completion was defined as when gaze position entered a 2° window around the saccade target (note that additional post-hoc analyses were conducted with alternative methods of defining saccade completion). After a variable post-saccadic delay, a large scene image (28° × 21°) was presented. In Experiment 1A, the scene image was presented either 5 ms or 500 ms after the recorded saccade offset, the 5 ms and 500 ms post-saccadic delay condition, respectively. The scene image was always presented for 50 ms, followed by a noise mask (500 ms). After the mask disappeared, subjects reported the category of scene image using a keyboard: S, D, F, J, K, and L. Correspondence between the six keys and the six scene categories were randomly assigned for each subject. Feedback for slow saccade reaction time was presented at the end of each trial if the saccade reaction time for the current trial was longer than 500 ms (“Eye movement too slow!”). Feedback for category reports was provided for 1000 ms only in practice trials (“Correct” or “Incorrect”). Subjects pressed the spacebar to continue to the next trial. Note that in the gaze-contingent design, the current trial was aborted and restarted after calibration if the subject failed to fixate on the initial fixation after more than 5 seconds after the onset of initial fixation, or failed to maintain fixation more than three times.

Experiment 1A included 24 practice trials using only full-spectrum (FS) scene images (see *Stimuli* section for details). In the first practice trial, images were presented for 150 ms, then linearly reduced to 29 ms over the course of 24 trials to familiarize participants with the task. During the main session, scene images were always presented for 50 ms and belonged to one of three spatial frequency conditions: full-spectrum (FS), high-spatial frequency (HSF), or low-spatial frequency (LSF). The experiment followed a 3 (Spatial Frequency: FS, HSF, LSF) × 2 (Post-Saccadic Delay: 5 ms, 500 ms) × 6 (Scene Category) design, with each condition repeated 10 times, resulting in 360 trials presented in random order across six blocks.

#### Experiment 1B

Experiment 1B was modified from Experiment 1A to examine a time course of post-saccadic scene processing by adding intermediate post-saccadic delay conditions, resulting in five post-saccadic delay conditions logarithmically spaced between 5 and 500 ms (5, 16, 50, 158, and 500 ms). Additionally, only LSF and HSF scene images, but not FS scene images, were used in both practice and main sessions to maximize number of trials for conditions of interests in a single session. Moreover, trials advanced automatically without requiring a spacebar press, and the saccade direction was always either horizontal or vertical, instead of diagonal, to ensure consistent saccade distance across trials. The main session followed a 2 (Spatial Frequency: LSF, HSF) × 5 (Post-Saccadic Delay: 5, 16, 50, 158, 500 ms) × 6 (Scene Category) design, with each condition repeated 10 times, totaling 600 trials, presented in random order across 10 blocks.

### Stimuli

Scene images and MATLAB code to filter the spatial frequency of scene images were modified from Perfetto et al. (2020). To create LSF and HSF scene images, grayscaled scene images were deconstructed using a two-dimensional Fast Fourier Transformation (FFT) and filtered with a low-pass (< 1 cycles per degree; cpd) or high-pass (> 6 cpd) SF filter with a 2^nd^-order Butterworth shaped boundary. The choice of the 2^nd^-order Butterworth filter and frequency cutoffs was based on Perfetto et al. (2020), where the scene categorization accuracy was comparable between HSF and LSF scene images (Experiment 1B; *t*(17) = 0.034; *p* = 0.97). The unfiltered full-spectrum image, HSF, and LSF version of a single image were jointly contrast-normalized (Figure 1B). The scene image was presented in size of 28° ×21° in the behavioral experiments.

We created 50 noise images to be randomly presented as a mask in each trial. To create noise images that contain low-level visual properties of scene images without identifiable category-specific features, we first calculated the average amplitude spectrum across all 432 scene images. Then, we performed inverse FFT using average amplitude and 50 random phase matrices to create 50 mask images. The 50 mask images were jointly contrast-normalized and rescaled to a range between 0.2 and 0.8.

There were 72 exemplar scene images (800 × 600 pixels) for each of the six scene categories (e.g., beach, city, forest, highway, mountain, office; Figure 1B). In Experiment 1A, scene images for each scene category (72 scenes) were divided into 6 groups. 1 group of scene images (12 scenes) was presented during the practice session, while the rest of the 5 groups (60 scenes) were used in the main session.

During the main session, each image was shown only once. Potentially, scene images could be identified more easily when filtered with either low or high spatial frequency bands. For example, forest scene images full of trees should be easier to recognize when filtered with high spatial frequency because of prevailing high spatial frequency information in vertical orientation (i.e., trees). To address such concern, we counterbalanced three spatial frequency conditions across the scene images presented during the main session between subjects. Specifically, for every group of three subjects, we used the same 12 scenes for practice trials, and the 60 scene images of the main session were divided into three sets of 20 images to be assigned to three spatial frequency conditions. In Experiment 1B, a scene image was randomly selected for each trial out of 72 exemplars from a given scene category.

### Apparatus

Experiments 1A and 1B were performed using MATLAB (The MathWorks, Natick, MA) with the Psychophysics Toolbox (Version 3 extension; Brainard, 1997; Kleiner, 2007; Pelli, 1997). Either the left or right eye position was monitored with the sampling rate of 1000 Hz using EyeLink 1000 eye-tracking system mounted on the desk, controlled by the Eyelink MATLAB Toolbox (Cornelissen et al., 2002). The eye-tracking system was calibrated using a nine-point grid method, at the beginning of the experiment and between trials if necessary.

Experiments were performed on a desk-top setting with 24.5-inch LCD monitor (ASUS ROG PG258Q) connected to NVIDIA GeForce RTX 2060, running at a 240 Hz refresh rate with a resolution of 1920 ×1080 pixels, located 63 cm in front of the participants (39 pixels per degree visual angle). Stimuli were presented above gray background (114 cd/m^2^) throughout the experiment.

### Behavioral data analysis

To examine post-saccadic processing of semantic category information, it is critical to validate actual stimuli duration and stimuli onset latency relative to the detection of saccade offset through post-hoc analysis of eye-tracking data. We confirmed that errors in these temporal manipulations were negligible. Experiment 1A showed scene duration close to 50 ms (mean = 48.47, sd = 1.81) and precise post-saccadic delay in 5 ms (mean = 3.73, sd = 1.76), and 500 ms post-saccadic delay trials (mean = 500.15, sd = 1.50) for all subjects. Experiment 1B showed accurate scene image duration (mean = 48.51, sd = 1.56), post-saccadic delay in 5 ms (mean = 3.51, sd = 1.50), 16 ms (mean = 14.52, sd = 1.46), 50 ms (mean = 48.84, sd = 1.54), 158 ms (mean = 156.71, sd = 1.54), and 500 ms post-saccadic delay trials (mean = 498.57, sd = 1.51).

In Experiment 1A, scene categorization accuracy for FS scene images was used to exclude subjects. The main analysis used data from LSF and HSF condition trials; scene categorization accuracies were compared by conducting 2 (Post-saccadic delay) × 2 (Spatial frequency) repeated measures ANOVA to examine the effect of saccade on subsequent scene perception and the modulation effect of spatial frequency. As pre-registered, in Experiment 1B, we conducted a 5 (Post-saccadic delay) × 2 (Spatial frequency) repeated measures ANOVA to test the main effect of post-saccadic delay, and how the spatial frequency condition modulates the main effect of post-saccadic delay. The spatial frequency condition was collapsed if the interaction term was not significant. If we find a significant main effect of post-saccadic delays on scene categorization accuracy, but not the interaction effect, we preregistered to perform post hoc *t*-tests after collapsing the spatial frequency condition. Specifically, scene categorization accuracy in the 500 ms post-saccadic delay condition (baseline) was compared with the remaining four shorter post-saccadic delay conditions (5, 16, 50, 158 ms). For each pairwise t-test, we used a critical alpha value of .0125, accounting for the number of *t*-tests performed (i.e., Bonferroni correction). Lower categorization accuracy compared to 500 ms delay condition will indicate disrupted processing of semantic scene category information.

In addition to the pre-registered analysis, we tested whether decreased categorization accuracy in shorter post-saccadic delay conditions is attributed to residual eye movement after saccade offset. First, we calculated eye movement velocity (°/sec) at each time point using a 10 ms sliding window for each trial. Then, we excluded trials in which eye movement velocity at scene onset was faster than 25 °/sec.

Because of a small number of remaining trials with shorter post-saccadic delays, we compared scene categorization accuracy between short post-saccadic delay trials (5 and 16 ms) and long post-saccadic delay trials (50, 158, and 500 ms). Second, to generalize the result with different saccade detection algorithms, we calculated post-saccadic delay for each trial based on the built-in online parsing system of Eyelink 1000 that incorporates eye movement velocity and acceleration rate to define saccade onset and offset. Using re-calculated post-saccadic delays, we separated trials into three post-saccadic delay groups (0-16 ms, 16-250 ms, and 250-1000 ms) and compared mean categorization accuracy.

### Experiment 2

#### Participants

17 subjects (14 women, 3 men, 0 nonbinary; age_mean_ = 23.56, age_std_ = 3.84) with normal or corrected-to-normal vision completed Experiment 2 (fMRI study). The sample size (N = 17) for Experiment 2 was determined through a priori power analysis using G*Power version 3.1.9.6 (Faul et al., 2009) based on a previous study (Berman et al., 2017), in which scene category decoding accuracy for high spatial frequency scene images in PPA was significantly above chance level (*t*(9) = 2.68, *p* = .025, *d* = 0.85). The power analysis estimated a required sample size of 17 for this effect size with a significance criterion (α) of 0.05 and a power of 0.9. All participants provided informed consent and were pre-screened for MRI eligibility. The study protocol was approved by the Ohio State University Biomedical Sciences institutional review board.

#### Experiment design

Subjects completed a 0.5-hour pre-scan session outside of the fMRI scanner and a 2-hour scan session in an fMRI scanner on different days. In both sessions, subjects were asked to follow a fixation dot and perform a 1-back task on sequentially presented scene images (Figure 4A). In each trial, the initial fixation dot was presented at one corner of an imaginary square (7° × 7°) for 1000 ms. Then, the initial fixation dot disappeared, and a saccade cue was presented at a new fixation location displaced horizontally or vertically by 7°, followed by the presentation of a large, full-field scene image for 100 ms. The task was to compare the scene image on the current trial to the one seen on the immediately prior trial (1-back task). Subjects were instructed to press a button only when a completely identical scene image was repeated, based on both content and spatial frequency, and to not press the button otherwise. Stimulus onset asynchrony (SOA) between trials was 4 seconds (50%), 6 seconds (33%), or 8 seconds (17%).

As a critical manipulation, we varied the timing of the saccade cue onset relative to scene onset across trials (Post-saccadic delay condition), such that the scene was presented after either a short (0-100 milliseconds) or long (400-600 milliseconds) post-saccadic delay. To achieve this, we used a different approach than the online gaze-contingent design employed in the behavioral experiments. In the fMRI experiment, the scene onsets had to be pre-determined and time-locked to the scanner’s repetition time (TR; 1,800 ms). Thus, we employed an approach where we measured the average saccadic reaction time for each subject in advance, and used this to individually adjust the time of saccade cue onset to maximize the number of trials where the scene images would be presented at the intended post-saccadic delays. We then performed post-hoc analyses of eye-tracking data for each subject to select trials where the scene image was actually presented within the intended short or long post-saccadic delay windows.

Specifically, we recorded saccade reaction times (SRT) - delay between saccade cue onset to saccade offset - from the pre-scan session (Figure 4B). From the SRT distribution, we identified a 100 ms time window encompassing most of the saccade reaction times (thick gray line in Figure 4B) and used the upper end of this window as the optimal saccade reaction time (optSRT; black arrow in Figure 4B).

During the scan session, the saccade cue was presented either optSRT or optSRT + 500 ms before the pre-determined time of scene onset, corresponding to the short and long post-saccadic delay conditions, respectively. For example, if participant’s optSRT was 250 ms and the scene image is scheduled to be presented 4,000 ms after the trial onset, the saccade cue appeared at either 3,750 ms (short delay) or 3,250 ms (long delay) after trial onset. With this approach, actual post-saccadic delays of individual subject followed a bimodal distribution, with peaks located between approximately 0–100 ms and 400–600 ms (Figure 4C). For each subject, trials which actually fell within the intended short or long post-saccadic delay windows were selected through post-hoc analyses of eye-tracking data to be included in the main analysis (Figure 4C colored portion of histograms; see Supplementary Figure 1 for individual subjects). Note that, due to individual variability of saccade onset latency, the number of included short and long delay condition trials differed between subjects (Supplementary Figure 1).

The scan session included 8 runs, each consisting of 108 trials with 12 repeated trials (repetition rate of 11.11%) where subjects had to press ‘1’; those trials were removed from further analysis. The rest of the 96 non-repeated trials comprised of 3 repetitions for each combination of 2 post-saccadic delay conditions (short and long post-saccadic delay) × 4 scene categories (mountain, beach, highway, and city) × 4 SF conditions (LSF1, LSF2, HSF1, and HSF2). The pre-scan session included two practice runs identical to that of the scan session, except that visual feedback (300 ms) was provided during the first run of the pre-scan (green: correct/red: incorrect).

Additionally, the scan session included one functional localizer run to localize early visual cortex and scene-selective regions (see *Region-of-Interest selection* section for details). The functional localizer run included 11 blocks: four object blocks, four scene blocks, and three fixation blocks. The order of blocks was counterbalanced across participants. In each block, 20 images were presented sequentially at the screen center (17.42° ×17.42°) for 400 ms with 500 ms delay. Participants performed a 1-back task with a repetition rate of 10%.

### Stimuli

To optimize stimuli for our fMRI design, we made a few changes to the scene stimuli from Experiment 1. Most critically, we used four scene category conditions and four spatial frequency conditions, each with a hierarchical structure of superordinate and subordinate categories. The set of scene images for Experiment 2 (Figure 4D) contained four subordinate scene categories (beach, mountain, city, and highway), affiliated with two superordinate scene categories (nature and urban). Scene images were grayscaled and filtered to contain either low or high spatial frequency information. To match the hierarchical structure of the scene category manipulation, we used four spatial frequency ranges, two low spatial frequency conditions and two high spatial frequency conditions: LSF1 (< 0.8 cpd low-pass filter), LSF2 (< 1.6 cpd low-pass filter), HSF1 (4-5 cpd band-pass filter), and HSF2 (7-10 cpd band-pass filter). To enhance recognizability of the HSF-filtered scene images, which contained a narrower range of spatial frequency than those used in the behavioral experiments, we applied an additional image processing step and computed the absolute values of the HSF-filtered images, which produces more naturalistic and familiar images resembling line drawings (Perfetto et al., 2020, Experiment 3). The four SF-filtered images from a single scene were jointly contrast-normalized and equated with mean luminance. The scene image was presented in size of 23.23° ×17.42° in both the pre-scan and the scan session to fully cover the entire screen.

The fixation dot was configured as an inner circle (0.3° diameter) with a thick outline (0.2° width). During an inter-trial interval, the white inner circle was surrounded by a black outline. The fixation dot changed to a black circle with a white outline (i.e., initial fixation onset; Figure 4A) 1 second prior to saccade cue onset to encourage subjects to fixate on the dot.

### Apparatus

The pre-scan and scan session of Experiment 2 were both performed using MATLAB (The MathWorks, Natick, MA) with the Psychophysics Toolbox (Version 3 extension; Brainard, 1997; Kleiner, 2007; Pelli, 1997). The pre-scan session of Experiment 2 was performed at the same setting as Experiment 1.

The scan session of Experiment 2 was carried out in a Siemens Prisma 3-T MRI scanner with an integrated Total Imaging Matrix (TIM) system using a 32-channel phased array receiver head coil, located at the OSU Center for Cognitive and Behavioral Brain Imaging. Functional data were acquired using a T2-weighted gradient-echo sequence (repetition time = 1,800 ms, echo time = 28 ms, flip angle = 70°). We used multiband whole-brain coverage aligned to the AC–PC (72 slices, 2 × 2 × 2 mm voxel, 10% gap, multiband factor = 3). Before the functional scan, a T1-weighted magnetization-prepared rapid gradient echo anatomic scan at 1-mm^3^ resolution was collected. Visual stimuli were presented using a rear-projection screen powered by a 3-chip DLP projector with a refresh rate of 60 Hz and a spatial resolution of 1280 × 1024 pixels. Participants lay down in the scanner and viewed stimuli distanced 74 cm via a mirror tilted 45° above the head coil. The EyeLink1000 eye-tracking system was positioned to monitor the right eye through the mirror with the sampling rate of 1000 Hz. To prevent the head coil from blocking the view of the right eye, subjects were repositioned slightly to the right when necessary. The eye-tracking system was calibrated using a nine-point grid method at the beginning of the experiment and between runs if necessary.

### fMRI data analysis

#### Preprocessing

fMRI data obtained from the functional localizer runs and main task runs were both corrected for slice acquisition time and head motion, and registered into Talairach space (Talairach & Tournoux, 1988) using Brain Voyager QX (Brain Innovation Maastricht, The Netherlands; Goebel et al., 2006). Different pre-processing steps were applied for fMRI data from the functional localizer run and main task runs.

The fMRI data from the functional localizer run were pre-processed with temporal filtering (GLM Fourier, two cycles) and spatial smoothing using a 4-mm FWHM Gaussian kernel. A whole-brain random-effects GLM was then applied to estimate beta coefficients for fixation, scene, and object blocks.

For fMRI data from main task, we calculated single-trial bata estimates for each onset of scene images (i.e., 864 trials) using GLMsingle toolbox (Prince et al., 2022), which was developed to optimize the estimation of single-trial fMRI responses using advanced denoising techniques (Kay et al., 2013; Rokem & Kay, 2020). Spatial smoothing was not performed because we planned to run MVPA analysis. Moreover, no temporal filtering was applied before running GLMsingle because GLMsingle accounts for baseline signal drift within runs by incorporating polynomial regressors into the model (Kay et al., 2013; Prince et al., 2022). The design matrices used for the GLMsingle included 16 conditions (4 spatial frequency conditions × 4 scene category conditions), without the post-saccadic delay condition to avoid systematic differences in estimated trial-wise beta coefficients between short and long post-saccadic delay trials.

#### Region-of-Interest selection

We defined functional regions of interest (ROIs) for individual subjects using GLM contrasts between scene, object, and fixation blocks during the functional localizer run. As a primary scene-selective ROI, we localized the parahippocampal place area (PPA; R. Epstein & Kanwisher, 1998), *scenes > objects contrast* contrast. Clusters of voxels showing significant activation were selected in volume space, with thresholds adjusted individually across subjects (most subjects: *p* < 0.01, a few subjects: up to *p* < 0.035; Supplementary Figure 2).

For exploratory analysis motivated by the functional distinction of PPA along the anterior-posterior axis (Baldassano et al., 2016; Berman et al., 2017; Epstein & Baker, 2019; Steel et al., 2024), we further divided PPA into anterior PPA (aPPA) and posterior PPA (pPPA) for each participant to have an equal number of voxels between the two PPA subregions. We also defined additional scene-selective regions: the retrosplenial complex (RSC; R. A. Epstein, 2008; O’Craven & Kanwisher, 2000) and the occipital place area (OPA; Dilks et al., 2013; Nakamura, 2000) for supplemental analyses.

For low-level visual analyses, an early visual cortex (EVC) ROI was localized using an *all conditions* > *fixation* contrast (similar to Golomb & Kanwisher, 2012a). Specifically, we applied a *[scenes & objects] > fixation* contrast, with a variable threshold (*p* < 0.000001 to *p* < 0.005; Supplementary Figure 2) and knowledge of anatomical boundaries (Wandell et al., 2007) to localize a region roughly including V1-V3 for each subject.

#### MVPA analysis

We used multi-voxel pattern analysis (MVPA) to quantify semantic scene category information in PPA using representational similarity calculated from correlation matrices (Haxby et al., 2001; Golomb & Kanwisher, 2012a). Trials that met the eye-tracking inclusion criteria for one of the two delays were coded into 32 conditions (Figure 5A): 2 post-saccadic delay conditions (short and long) × 4 scene category conditions (beach, mountain, city, and highway) × 4 spatial frequency conditions (LSF1, LSF2, HSF1, and HSF2). We then calculated a 32 × 32 representational similarity matrix (RSM) using the split-half correlation method. First, we split the 8 runs into two groups of runs. Single-trial beta estimates were averaged across all trials with the same condition label within each group of runs. Then, for each group of runs separately, we normalized each voxel’s response by subtracting the beta coefficient averaged across conditions from the beta estimates of each condition. Lastly, the voxel-wise beta coefficients for each of the 32 conditions in one group of runs were correlated with each of the 32 conditions in the other group of runs, and Pearson’s r values were converted to z-scores using Fisher’s z transformation, generating a 32 × 32 correlation matrix for PPA (Figure 5B). All subsequent analyses were performed on the z-scored data.

Notably, when splitting data into two groups of runs, we first followed the conventional odd versus even runs split-half method (Haxby et al., 2001). However, because we excluded 1-back repeat trials and those with a post-saccadic delay outside the range of either the short or long post-saccadic delay, the number of trials remaining for each condition in each group of runs was not only different but also sometimes zero. If there was a condition in either group of runs with no included trials, we randomly re-divided the eight runs into two groups until there was at least one trial for every condition (Supplementary Figure 3). There were typically more trials in the long delay conditions compared to the short delay conditions. To account for the different number of trials between the short and long delay conditions, we performed a control analysis where we down-sampled the long delay trials to match the number of short delay trials for each condition per each group of run (e.g., equal number of trials between condition 1 and 17, or 2 and 18, etc. in Supplementary Figure 3). Nevertheless, the pattern of results remained the same (Supplementary Figure 4), and therefore, we focus on the results without down-sampling.

From the RSM, we quantified the amount of scene category information (Nature vs. Urban) in the short or long-delay trials, separately for LSF and HSF trials (Figure 5C). First, we divided the 32 × 32 correlation matrix into four 8 × 8 subsets, each corresponding to different post-saccadic delay conditions (Short vs. Long) and spatial frequency conditions (LSF vs. HSF). Then, we calculated the average representational similarity (average z-score) for the cells corresponding to the same and the different scene category pairs, and took their difference as an index for scene category representation. For example, to calculate scene category information (Nature vs. Urban) in the short post-saccadic delay HSF trials, we selected the subset of RSM cells corresponding to the short post-saccadic delay and HSF conditions (Figure 5C, third from the left). Then, we subtracted the average similarity between the *different* scene category pairs (Figure 5C; black cells) from the *same* scene category pairs (Figure 5C; white cells). If the voxel-wise response pattern is more similar (i.e., higher correlation) between conditions sharing the *same* scene category than for those with *different* scene categories (significantly positive value after subtraction), then that indicates the neural activity pattern in PPA contains a representation of scene category information (Haxby et al., 2001). In the main text, we report the results of our primary analyses on the PPA, but in the supplement include exploratory analyses investigating scene category information in other scene-selective brain regions (i.e., RSC, OPA; Supplementary Figure 5).

We also examined how the representation of a low-level visual feature, spatial frequency, is influenced post-saccadically, by performing analogous MVPA analysis quantifying spatial frequency information, focusing on early visual cortex (EVC).

Note that for our main analyses, we focused primarily on superordinate-level scene category (Nature vs. Urban) and spatial frequency (LSF vs. HSF), combining across subordinate-level scene category and spatial frequency information to improve power and MVPA performance. Nevertheless, we observed similar results when decoding subordinate-level scene category (Beach vs. Mountain vs. City vs. Highway). Additionally, we examined subordinate-level spatial frequency information in EVC, separately for low (LSF1 vs. LSF2) and high (HSF1 vs. HSF2) spatial frequency bands (Supplementary Figure 6). We also report analyses separating the RSM only by delay (16 x 16 cell RSMs) to explore effects of post-saccadic delay on scene category information regardless of spatial frequency.

For all statistical testing of results, we used both frequentist and Bayesian approaches using JASP software (Version 0.14; JASP, 2017).

## Acknowledgements

This work was supported by the National Institute of Health [NIH R01-EY025648 (JG)]; the National Science Foundation [NSF 1848939 (JG)].

## Supplemental Materials

**Supplementary Figure 1.**
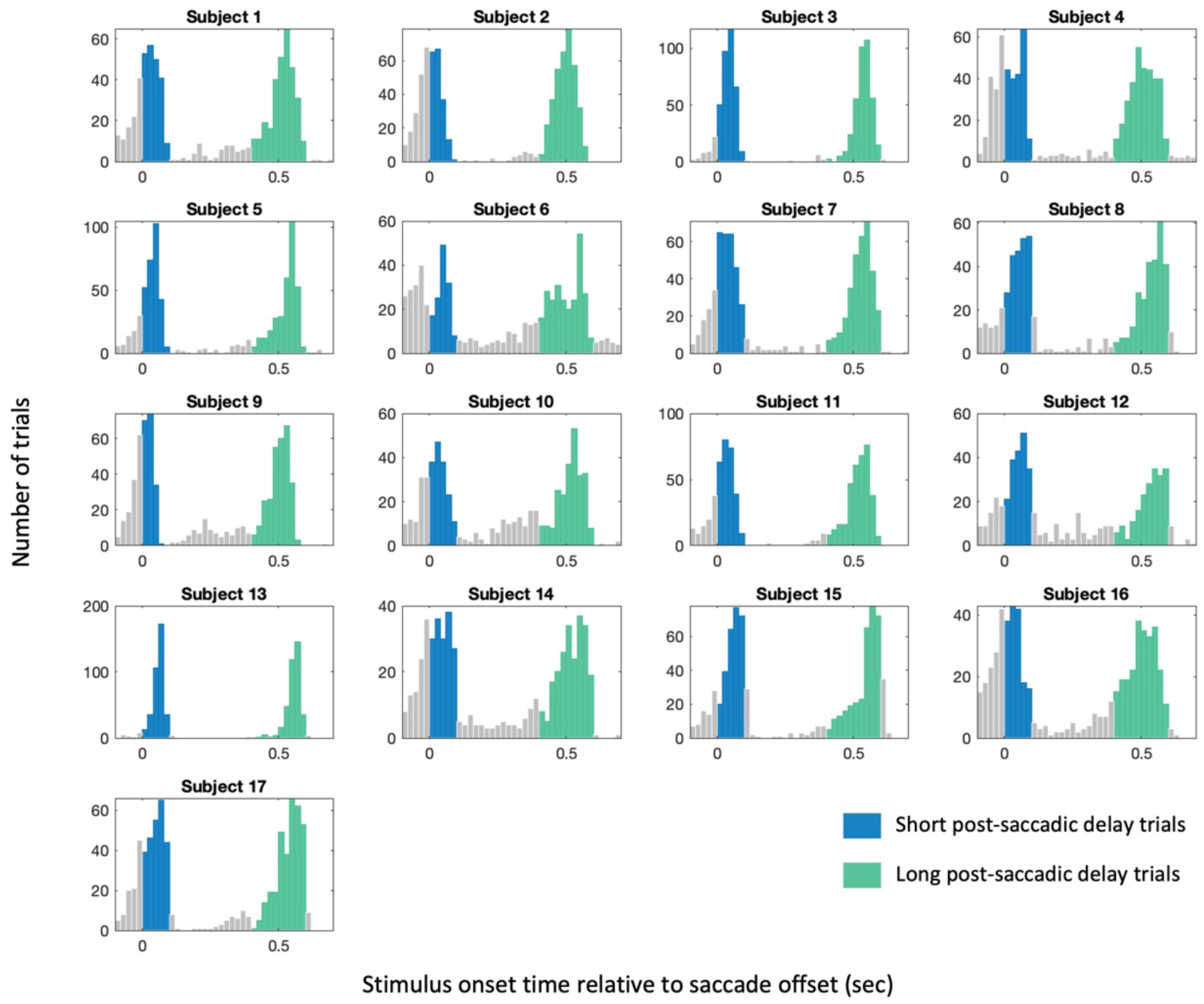
Distribution of stimulus onset times relative to saccade offset for individual subjects (N=17) who completed Experiment 2. Trials with stimulus onset time relative to saccade offset within a 0-100 ms window were labeled as short post-saccadic delay trials (blue), while those within a 400-600 ms window were labeled as long post-saccadic delay trials (green).

**Supplementary Figure 2.**
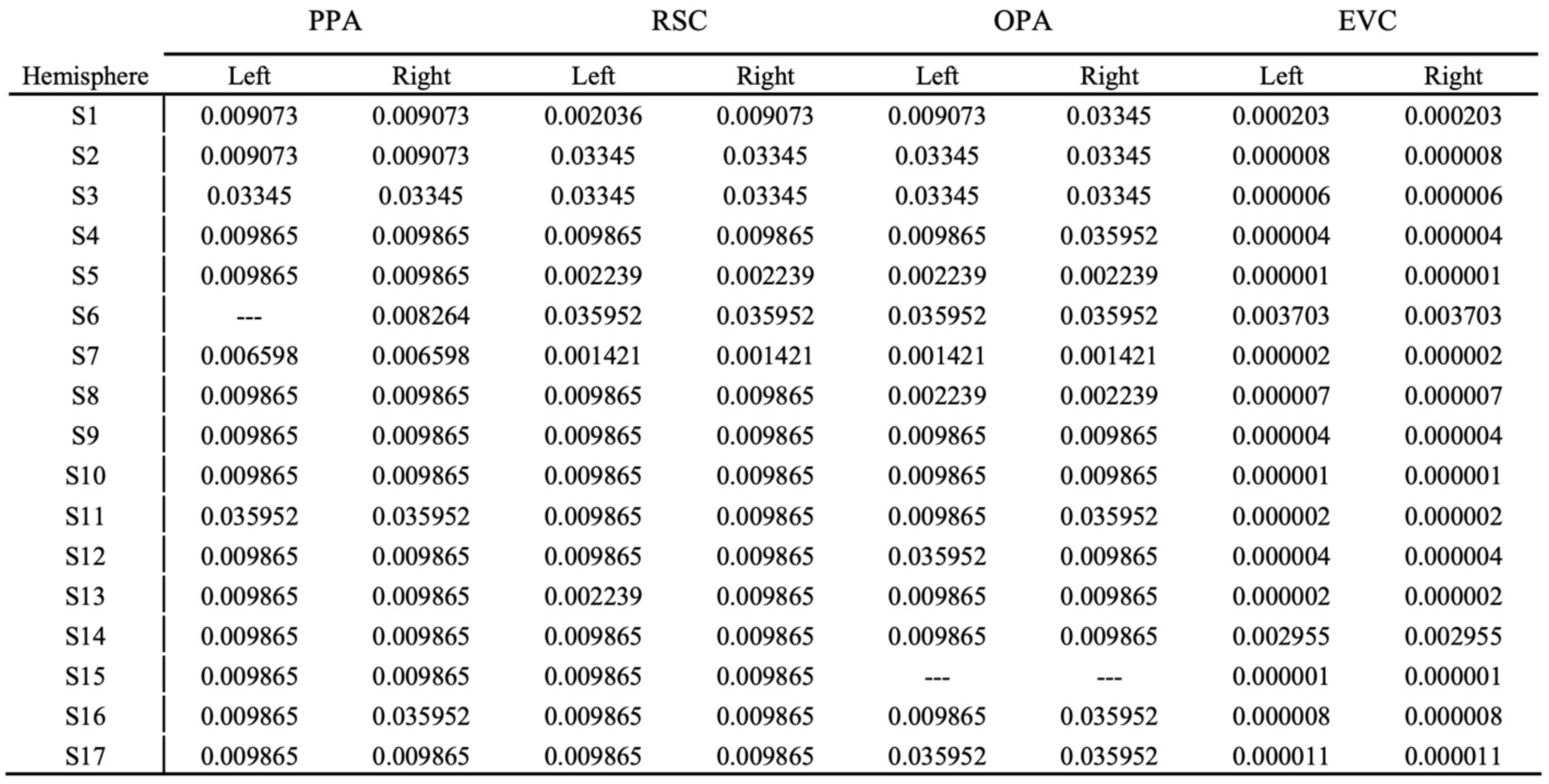
*p*-values used to select a cluster of voxels defined as regions of interest in Experiment 2. Functional ROIs were defined individually for each subject using GLM contrasts between scene, object, and fixation blocks from the functional localizer run. As the primary scene-selective ROI, we identified the parahippocampal place area (PPA; Epstein & Kanwisher, 1998) using the *scenes > objects* contrast. Additional ROIs included the retrosplenial complex (RSC; Epstein, 2008; O’Craven & Kanwisher, 2000) and the occipital place area (OPA; Dilks et al., 2013; Nakamura, 2000), which were used in supplemental analyses. Voxel clusters showing significant activation were selected in volume space, with subject-specific thresholds (most subjects:*p <* 0.01; a few subjects: up *top* < 0.035). For Subject 6, no significant cluster near the left PPA could be identified even at a relaxed threshold of/? < 0.05; therefore, only the right PPA was included for that subject. For Subject 15, OPA could not be localized in either hemisphere, and the subject was excluded from OPA analyses.

**Supplementary Figure 3.**
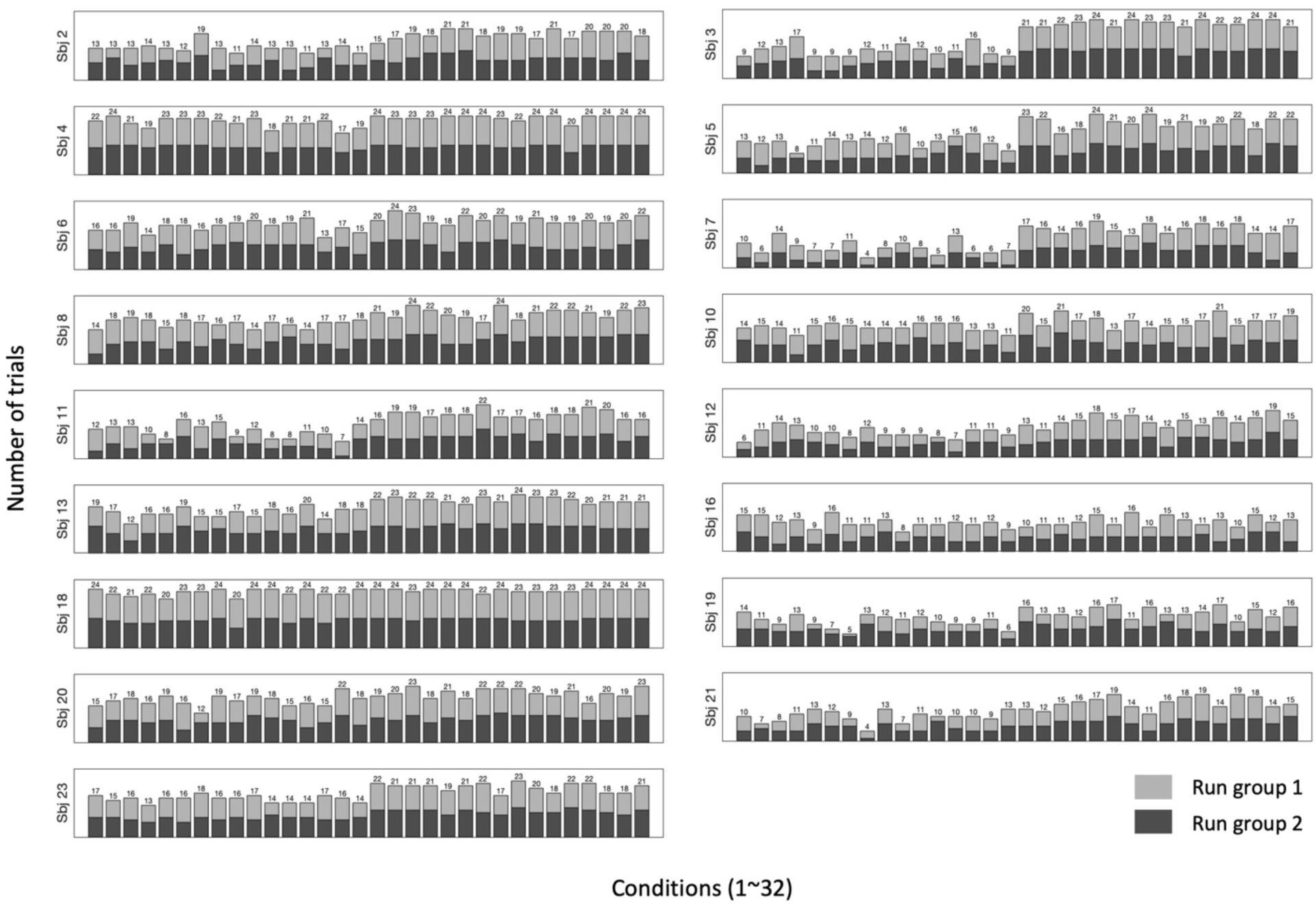
Number of trials included in each condition to construct representational similarity matrix in Experiment 2. Two differently shaded bars indicate two groups of runs.

**Supplementary Figure 4.**
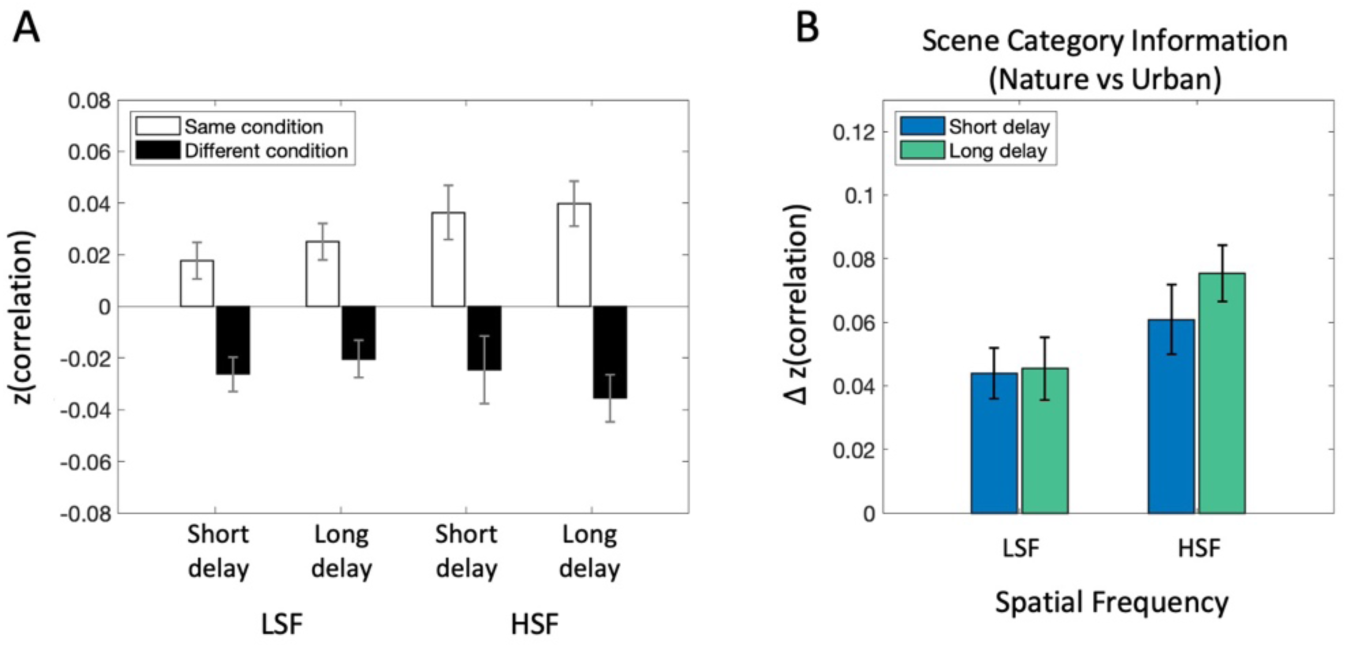
fMRI MVPA analysis results in PPA matched in the number of trials between the short and long post-saccadic delay condition. For the main analysis, we used a broader time window for long post-saccadic trials (400-600 ms) compared to the short delay trials (0-100 ms). Thus, the number of trials used to construct RSM was smaller for short than the long post-saccadic delay conditions (Supplementary Figure 3). Since the number of trials could influence MVPA results, we conducted a control analysis where we matched the number of trials between short and long post-saccadic delay trials. (A) Averaged correlation between the same scene category pairs (black bars) and different category pairs (white bars), demonstrating more similar neural activity patterns between same scene category trials than different scene category trials. Statistically, we found a significantly larger averaged correlation between same scene category pairs compared to different scene category pairs (*ps* < .018, *BF*_10_s > 3.26). (B) Scene category representation in PPA, obtained by subtracting average correlation between different scene category pairs from same category pairs, separately for spatial frequency conditions (LSF vs. Long) and post-saccadic delay conditions (Short vs. Long). 2 (Post-saccadic delay condition) × 2 (Spatial frequency condition) repeated measures ANOVA revealed significant main effect of post-saccadic delay (F(l, 16) = *6.22, p* = .024, 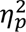 *=* .28, *BF*_incl_= 0.68), without main effect of spatial frequency (F(l,16) = 4.23, *p* = .056, 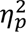 = .21, *BF_mü_ =* 1.78) nor the interaction effect (F(l,16) = 0.83, *p* = .377, 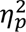 = .049, *BF*_incl_ = 0.51). The consistent pattern of results further recapitulates the post-saccadic reduction of scene category information represented in PPA. Error bars indicate within-subject standard error.

**Supplementary Figure 5.**
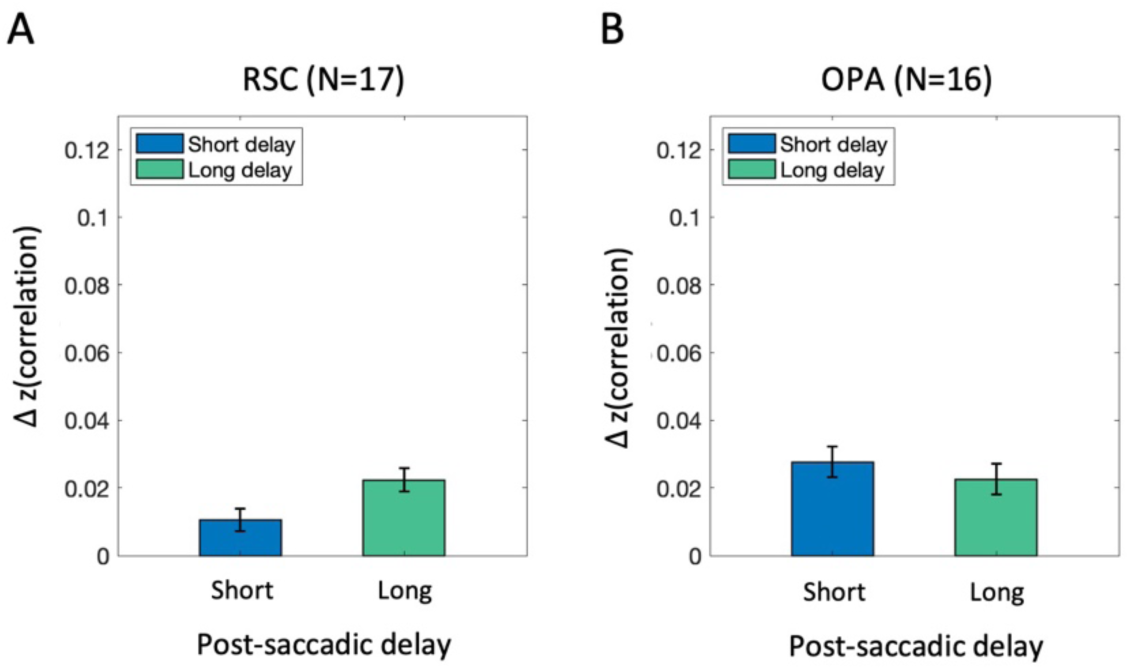
MVPA analysis results in other scene-selective regions (i.e., RSC and OPA) that were both anatomically and functionally distinct from each other (Epstein & Baker, 2019; Epstein & Kanwisher, 1998; Wolbers & Biichel, 2005).

We found no significant effect of post-saccadic delay condition on scene category representation both in RSC (A; t(l6) = −1.77,p = .10, *d* = −0.43, *BF_10_* = 0.90) and OPA (B; t(l5) = .57,p = .580, *d* = 0.141, *BF_10_* = 0.29). Notably, no significant effect ofpost-saccadic delay may be in part due to the lack of power to capture semantic category information represented in these scene-selective regions. In RSC, there was a significant scene category information in long delay trials (t(16) = 3.86, *p* = .001, *d* = 0.94, *BF_10_* = 28.67), but not in short delay trials *(t(l* 6) = 1.23,p = .237, *d* = 0.30, *BF_10_* = 0.48). OPA showed significant scene category information in short delay trials (t(15) = 2.85, *p* = .012, *d* = 0.71, *BF_10_* =4.64), but not in the long delay trials (t(15) = 1.97,p = .067, *d=* 0.49, *BF_10_* = 1.20). Error bars indicate within-subject standard error.

Given previous findings showing that spatial frequency manipulations specifically affect scene processing in PPA (Musel et al., 2014; Berman et al., 2017), the current study focused primarily on the PPA in the ventral visual stream as a locus of scene processing. However, RSC (R. A. Epstein, 2008; O’Craven & Kanwisher, 2000) and OPA (Dilks et al., 2013; Nakamura, 2000) also play crucial roles in scene processing and are functionally connected (Baldassano et al., 2016). While we could functionally localize PPA, RSC, and OPA as individual ROIs, MVPA analysis did not yield significant scene category representation in RSC and OPA, potentially due to the insufficient presentation time of scene image (100 ms) and extensive preprocessing applied to scene images (e.g., gray-scaling, frequency filtering, etc.). While whether RSC or OPA would exhibit a similar pattern of disrupted post-saccadic scene information as PPA remains unclear, known functional distinctions between scene-selective regions (Epstein & Kanwisher, 1998; Epstein & Baker, 2019; Kauffmann et al., 2015; Wolbers & Biichel, 2005) imply a distinct effect of saccades between scene-selective regions. For instance, PPA and OPA may be more susceptible to saccade-induced disruption compared to RSC, since RSC is involved in the mnemonic aspect of scene processing, whereas PPA and OPA are more perceptually oriented (Epstein et al., 2007; Wolbers & Biichel, 2005).

**Supplementary Figure 6.**
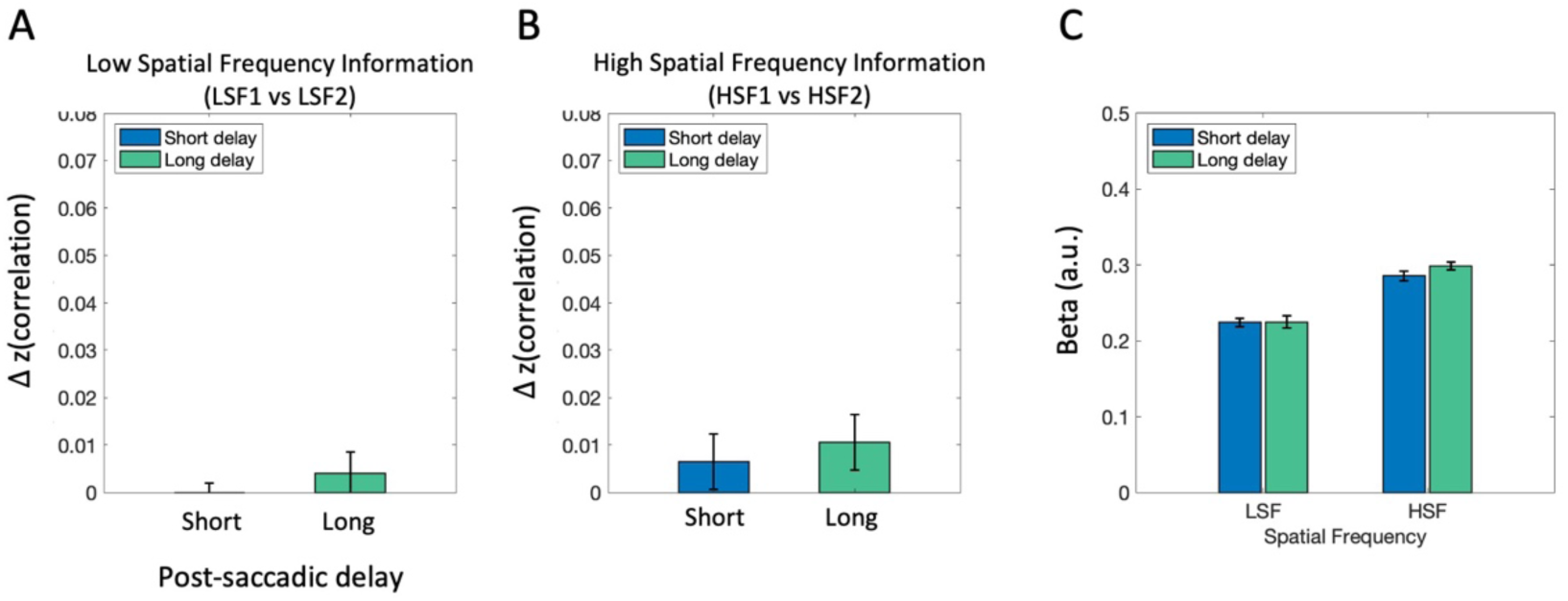
fMRI analysis results in EVC, broken down to subordinate spatial frequency information: low (LSF1 vs. LSF2) or high (HSF1 vs. HSF2) spatial frequency. Neural representation of spatial frequency was compared between the short and long post-saccadic conditions, separately for (A) low (LSF1 vs LSF2) and (B) high spatial frequency bands (HSF1 vs HSF2). We found no significant spatial frequency information for both short and long post-saccadic delay trials in the LSF condition (*ps* > .544, *BF*_10_*<* 0.295; Figure A) and the HSF condition (*ps* > .229, *BF*_10_*<* 0.49). (C) Likewise, univariate activation was compared across conditions, which showed Additionally, 2 × 2 ANOVA performed on univariate activation showed significantly stronger activation in EVC for HSF images compared to LSF scene images (F(l, 16) = 67.36, *p <* .001, 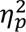 *=* .81, *BF*_incl_ *=* 1.52e+9), without significant interaction (F(l, 16) = 1.11, *p* = .309, 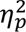 *=* .065, *BF*_incl_ *=* 0.45) nor main effect of post-saccadic delay (F(l, 16) = 75, *p* = .403, 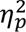 *=* .044, *BF*_incl_ = 0.34). Error bars indicate within-subject standard error.

