## Supplemental Materials for "Post-Saccadic Disruption of Semantic Category Information in Naturalistic Scenes"

### Supplementary Materials

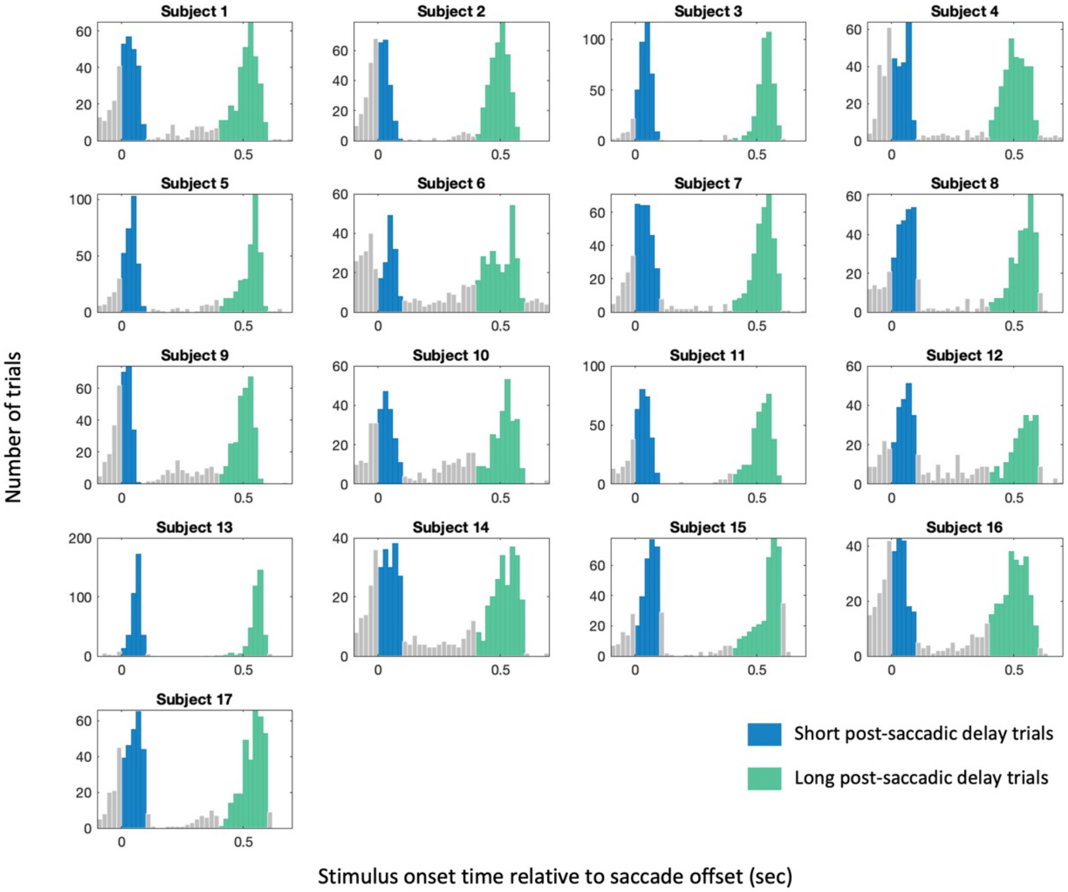


*Supplementary Figure 1.* Distribution of stimulus onset times relative to saccade offset for individual subjects (N=17) who completed Experiment 2. Trials with stimulus onset time relative to saccade offset within a 0-100 ms window were labeled as short post-saccadic delay trials (blue), while those within a 400-600 ms window were labeled as long post-saccadic delay trials (green).

| | PPA | | RSC | | OPA | | EVC | |
|---|---|---|---|---|---|---|---|---|
| Hemisphere | Left | Right | Left | Right | Left | Right | Left | Right |
| S1 | 0.009073 | 0.009073 | 0.002036 | 0.009073 | 0.009073 | 0.03345 | 0.000203 | 0.000203 |
| S2 | 0.009073 | 0.009073 | 0.03345 | 0.03345 | 0.03345 | 0.03345 | 0.000008 | 0.000008 |
| S3 | 0.03345 | 0.03345 | 0.03345 | 0.03345 | 0.03345 | 0.03345 | 0.000006 | 0.000006 |
| S4 | 0.009865 | 0.009865 | 0.009865 | 0.009865 | 0.009865 | 0.035952 | 0.000004 | 0.000004 |
| S5 | 0.009865 | 0.009865 | 0.002239 | 0.002239 | 0.002239 | 0.002239 | 0.000001 | 0.000001 |
| S6 | --- | 0.008264 | 0.035952 | 0.035952 | 0.035952 | 0.035952 | 0.003703 | 0.003703 |
| S7 | 0.006598 | 0.006598 | 0.001421 | 0.001421 | 0.001421 | 0.001421 | 0.000002 | 0.000002 |
| S8 | 0.009865 | 0.009865 | 0.009865 | 0.009865 | 0.002239 | 0.002239 | 0.000007 | 0.000007 |
| S9 | 0.009865 | 0.009865 | 0.009865 | 0.009865 | 0.009865 | 0.009865 | 0.000004 | 0.000004 |
| S10 | 0.009865 | 0.009865 | 0.009865 | 0.009865 | 0.009865 | 0.009865 | 0.000001 | 0.000001 |
| S11 | 0.035952 | 0.035952 | 0.009865 | 0.009865 | 0.009865 | 0.035952 | 0.000002 | 0.000002 |
| S12 | 0.009865 | 0.009865 | 0.009865 | 0.009865 | 0.035952 | 0.009865 | 0.000004 | 0.000004 |
| S13 | 0.009865 | 0.009865 | 0.002239 | 0.009865 | 0.009865 | 0.009865 | 0.000002 | 0.000002 |
| S14 | 0.009865 | 0.009865 | 0.009865 | 0.009865 | 0.009865 | 0.009865 | 0.002955 | 0.002955 |
| S15 | 0.009865 | 0.009865 | 0.009865 | 0.009865 | --- | --- | 0.000001 | 0.000001 |
| S16 | 0.009865 | 0.035952 | 0.009865 | 0.009865 | 0.009865 | 0.035952 | 0.000008 | 0.000008 |
| S17 | 0.009865 | 0.009865 | 0.009865 | 0.009865 | 0.035952 | 0.035952 | 0.000011 | 0.000011 |

*Supplementary Figure 2. p*-values used to select a cluster of voxels defined as regions of interest in Experiment 2. Functional ROIs were defined individually for each subject using GLM contrasts between scene, object, and fixation blocks from the functional localizer run. As the primary scene-selective ROI, we identified the parahippocampal place area (PPA; Epstein & Kanwisher, 1998) using the *scenes* > *objects* contrast. Additional ROIs included the retrosplenial complex (RSC; Epstein, 2008; O'Craven & Kanwisher, 2000) and the occipital place area (OPA; Dilks et al., 2013; Nakamura, 2000), which were used in supplemental analyses. Voxel clusters showing significant activation were selected in volume space, with subject-specific thresholds (most subjects: $p < 0.01$; a few subjects: up to $p < 0.035$). For Subject 6, no significant cluster near the left PPA could be identified even at a relaxed threshold of $p < 0.05$; therefore, only the right PPA was included for that subject. For Subject 15, OPA could not be localized in either hemisphere, and the subject was excluded from OPA analyses.

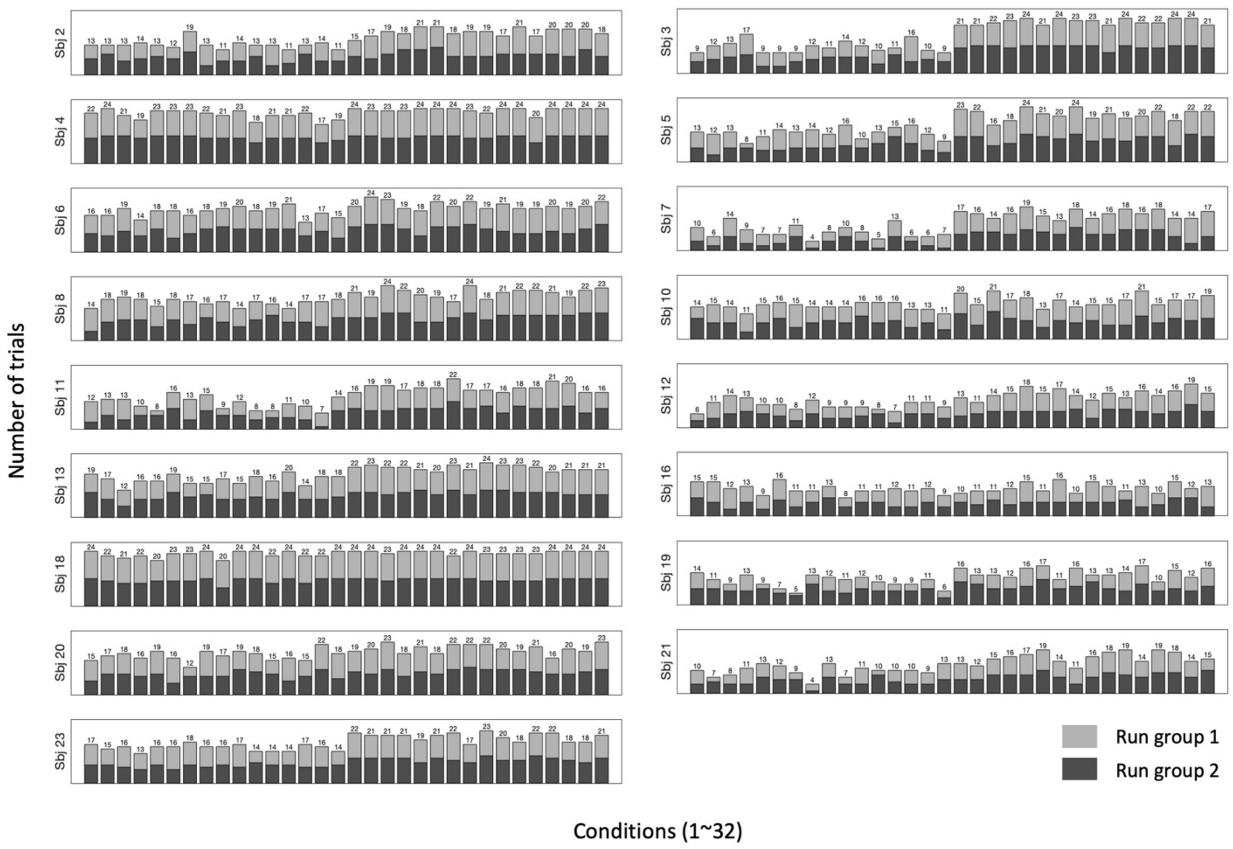


*Supplementary Figure 3*. Number of trials included in each condition to construct representational similarity matrix in Experiment 2. Two differently shaded bars indicate two groups of runs.

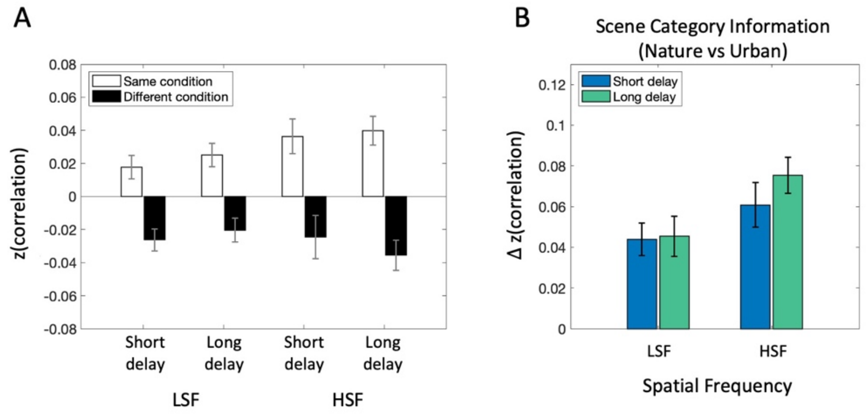


*Supplementary Figure 4.* fMRI MVPA analysis results in PPA matched in the number of trials between the short and long post-saccadic delay condition.

For the main analysis, we used a broader time window for long post-saccadic trials (400-600 ms) compared to the short delay trials (0-100 ms). Thus, the number of trials used to construct RSM was smaller for short than the long post-saccadic delay conditions (Supplementary Figure 3). Since the number of trials could influence MVPA results, we conducted a control analysis where we matched the number of trials between short and long post-saccadic delay trials.

(A) Averaged correlation between the same scene category pairs (black bars) and different category pairs (white bars), demonstrating more similar neural activity patterns between same scene category trials than different scene category trials. Statistically, we found a significantly larger averaged correlation between same scene category pairs compared to different scene category pairs ($p$s < .018, $BF_{10}$s > 3.26).

(B) Scene category representation in PPA, obtained by subtracting average correlation between different scene category pairs from same category pairs, separately for spatial frequency conditions (LSF vs. Long) and post-saccadic delay conditions (Short vs. Long). 2 (Post-saccadic delay condition) × 2 (Spatial frequency condition) repeated measures ANOVA revealed significant main effect of post-saccadic delay ($F(1,16) = 6.22$, $p = .024$, $\eta_p^2 = .28$, $BF_{incl} = 0.68$), without main effect of spatial frequency ($F(1,16) = 4.23$, $p = .056$, $\eta_p^2 = .21$, $BF_{incl} = 1.78$) nor the interaction effect ($F(1,16) = 0.83$, $p = .377$, $\eta_p^2 = .049$, $BF_{incl} = 0.51$). The consistent pattern of results further recapitulates the post-saccadic reduction of scene category information represented in PPA. Error bars indicate within-subject standard error.

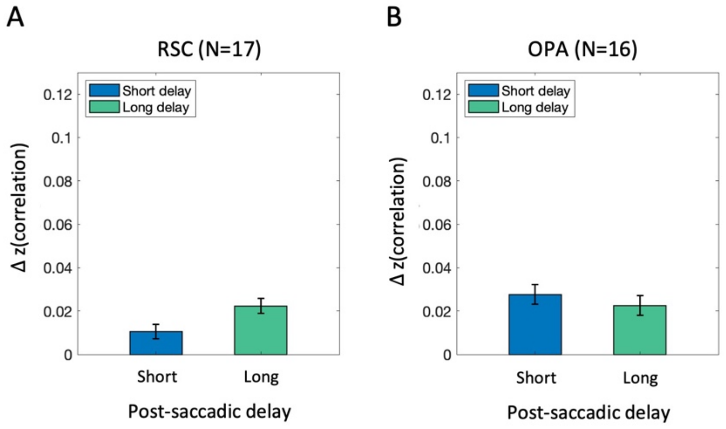


*Supplementary Figure 5.* MVPA analysis results in other scene-selective regions (i.e., RSC and OPA) that were both anatomically and functionally distinct from each other (Epstein & Baker, 2019; Epstein & Kanwisher, 1998; Wolbers & Büchel, 2005).

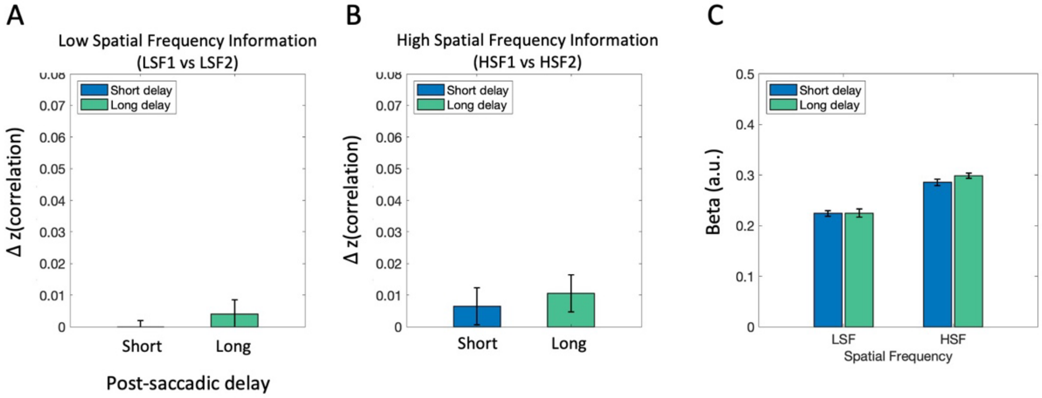


*Supplementary Figure 6.* fMRI analysis results in EVC, broken down to subordinate spatial frequency information: low (LSF1 vs. LSF2) or high (HSF1 vs. HSF2) spatial frequency. Neural representation of spatial frequency was compared between the short and long post-saccadic conditions, separately for (A) low (LSF1 vs LSF2) and (B) high spatial frequency bands (HSF1 vs HSF2). We found no significant spatial frequency information for both short and long post-saccadic delay trials in the LSF condition ($p$s > .544, $BF_{10} < 0.295$; Figure A) and the HSF condition ($p$s > .229, $BF_{10} < 0.49$). (C) Likewise, univariate activation was compared across conditions, which showed Additionally, 2 × 2 ANOVA performed on univariate activation showed significantly stronger activation in EVC for HSF images compared to LSF scene images (F(1,16) = 67.36, $p < .001$, $\eta_p^2 = .81$, $BF_{\text{incl}} = 1.52e+9$), without significant interaction (F(1,16) = 1.11, $p = .309$, $\eta_p^2 = .065$, $BF_{\text{incl}} = 0.45$) nor main effect of post-saccadic delay (F(1,16) = 75, $p = .403$, $\eta_p^2 = .044$, $BF_{\text{incl}} = 0.34$). Error bars indicate within-subject standard error.
